# Molecular mechanism of action of a blood brain barrier shuttle antibody

**DOI:** 10.64898/2026.06.30.735565

**Authors:** Sulin Liu, Ollie E King, Ayla A Wahid, Chuhan Shang, Luke A Pattison, Tomas Malinauskas, Eduardo A Santander, Stephanie A Nestorow, Wan-Na Chen, Vasiliki Mavridou, Jacob I. Browne, Ewan St. J Smith, John E Linley, Carl I Webster, Paul S Miller

## Abstract

The transferrin receptor has emerged as a prime target for transcytosis of antibody shuttles from the bloodstream into the brain parenchyma to deliver therapeutic payloads that treat neurological disorders. However, how the transferrin receptor-antibody binding mode impacts avidity, degradation, pH sensitivity and delivery remains underexplored. To address this, we determined the cryo-EM structure of mouse transferrin receptor 1 bound to the model brain shuttle antibody 8D3. In combination with cell binding and localisation studies we show that 8D3 can structurally support small-scale inter-receptor cross-linking, such as in self-contained pairs, that cause avidity but do not induce receptor redistribution or degradation. The structure can also explain pH-dependent binding modes, and we show how some of these antibody variants with graded sensitivities regulate brain penetration *in vivo*. Overall, our study illuminates how distinct molecular features of antibody binding impact transferrin receptor behaviour and brain delivery to inform on future shuttle design.

## INTRODUCTION

Systemic delivery of protein therapeutics to the brain is restricted by the blood-brain barrier (BBB), which contains an impermeant lining of brain endothelial cells (BECs) interconnected by tight junctions^1^. The use of molecular ‘Trojan horses’ has emerged as a frontrunner strategy for central nervous system (CNS) protein delivery^2–6^, whereby shuttle antibodies incorporating a therapeutic cargo are transcytosed across the BEC lining via endogenous receptors ^7^. Although several receptors have been investigated^8–16^, transferrin receptor 1 (TfR1, also known as CD71), which is responsible for transferrin-iron complex delivery and certain forms of virus-mediated entry^17–19^, is the most studied and only clinically proven target ^2,4,20–31^. TfR1 is a homodimeric type II transmembrane protein with each ectodomain monomer comprising helical, protease-like and apical subdomains^32^.

Delivery of antibody shuttles and cargo across the BBB is affected by both affinity and valency, with low-digit nanomolar affinity limiting release into the parenchyma^21,22,26,29,30,33^, and bivalent binding to TfR1 resulting in clustering, lysosomal targeting and degradation^21,34,35^. Antibody pH sensitivity to reduce TfR1 binding at low pH in endosomal compartments during transcytosis has also been shown to enhance transcytosis in an *in vitro* BBB model^36^. So far though, understanding how molecular binding influences avidity, degradation, delivery and pH sensitivity, is limited due to the absence of publicly available TfR1 PDB structures bound by antibody complementarity determining regions (CDRs)^21,30^. Furthermore, the limited TfR1 structures from non-human species ^37^, including mouse, hinders next-generation *de novo* BBB shuttle design approaches ^38–40^ to deliver cross-species binders ^30^. Instead, translational studies rely on transgenic hTfR1 knock-in mice, which might exhibit unnatural TfR1 behaviour and require time-consuming crossbreeding to other genetic disease strains^41,42^.

Here, we determine the cryo-EM structure of a model BBB shuttle antibody, 8D3^22,26,29,33,43,44^, bound to the mouse (m) TfR1 ectodomain. In combination with cell binding and localisation studies we explore the structural consequences on affinity versus avidity and mTfR1 distribution, showing contrasting effects versus alternative brain shuttles^21,30,34,35^. A structure-guided combinatorial histidine library allowed us to systematically explore the relationship between pH-selective binders and *in vivo* brain delivery. This study helps us to understand how the molecular binding mode influences brain delivery and provides a basis for designing better shuttles in the future.

## Results

### 8D3 binding mode and basis of selectivity

To understand the molecular action of the brain shuttle 8D3 we determined the structure of mTfR1 ectodomain in complex with an 8D3 Fab by cryo-EM at 2.9Å global resolution (Extended Data Fig. 1; Supplementary Table 1). Previously determined hTfR1 complexes show two binder interaction hotspots: along the helical domain helix-helix groove – hemochromatosis protein^45^, Tf^46^, DNA aptamer^47^, cystine dense peptide brain shuttle^48^; or at the apical domain – Machupo virus GP1^49^, *Plasmodium vivax* invasion complex^50^, computationally designed 3DS18 protein shuttle^51^, Denali Fc brain shuttle^52^, Parvovirus B19^53^ (Extended Data Fig. 2A). 8D3 binds to the mTfR1 apical domain at the same loops targeted on hTfR1 (Fig. 1a-c and Extended Data Fig. 2A). The heavy chain variable domain (V_H_) comprises the majority of the 8D3-mTfR1 interface, buried surface 480Å^2^, versus the light chain variable domain (V_L_), 200Å^2^. This is reflected by all three V_H_ CDRs contacting the apical domain versus only V_L_ CDR3. Between them the four CDR binding loops contain five tyrosine residues that crown two mTfR1 apical domain loops, a main loop (206-212) and a minor loop (371-373), themselves rich in asparagine and glutamine residues (6 from 10 residues) (Fig. 1c-e). The majority of interface contacts stem from V_H_. These include H-bond contacts (∼ 2.5-3.3 Å) from V_H_ peptide backbone amides or carbonyls: CDR3 Y103 with mTfR1 N209; CDR1 G33 and CDR3 P99 with mTfR1 N211 (Fig. 1e,f). Also, key vdW contacts arise between the V_H_ CDR2 Y52 and Y53 aromatic side chains and the main apical loop V206 and L212 aliphatic side chains (Fig. 1f). In contrast, the only interactions from V_L_ CDR3 are putative vdW contacts from Y92 to minor loop Q371 and W96 to main loop N209 (Fig. 1e). Consistent with the close juxtaposition of the 8D3 Tyr residues to mTfR1, previous studies have shown that LC-Y92A and HC-Y52A, Y53A, Y103A single mutations all reduce affinity by at least 10-fold^33,54^, and HC-Y32A reduced affinity 2-fold^29^. We measured on-cell binding apparent affinity at 4 °C using an HEK293 stable cell line expressing mTfR1 for wild type (WT) 8D3 versus a double substitution LC-Q90A HC-Y103A (8D3-MT), and this reduced the apparent affinity by ∼100-fold (Fig. 1g), similar to a previous study^22^. Furthermore, alanine substitution to 6 of these residues together to generate 8D3-KO, eliminated binding up to 1 µM (Fig. 1g).

**Fig. 1.**
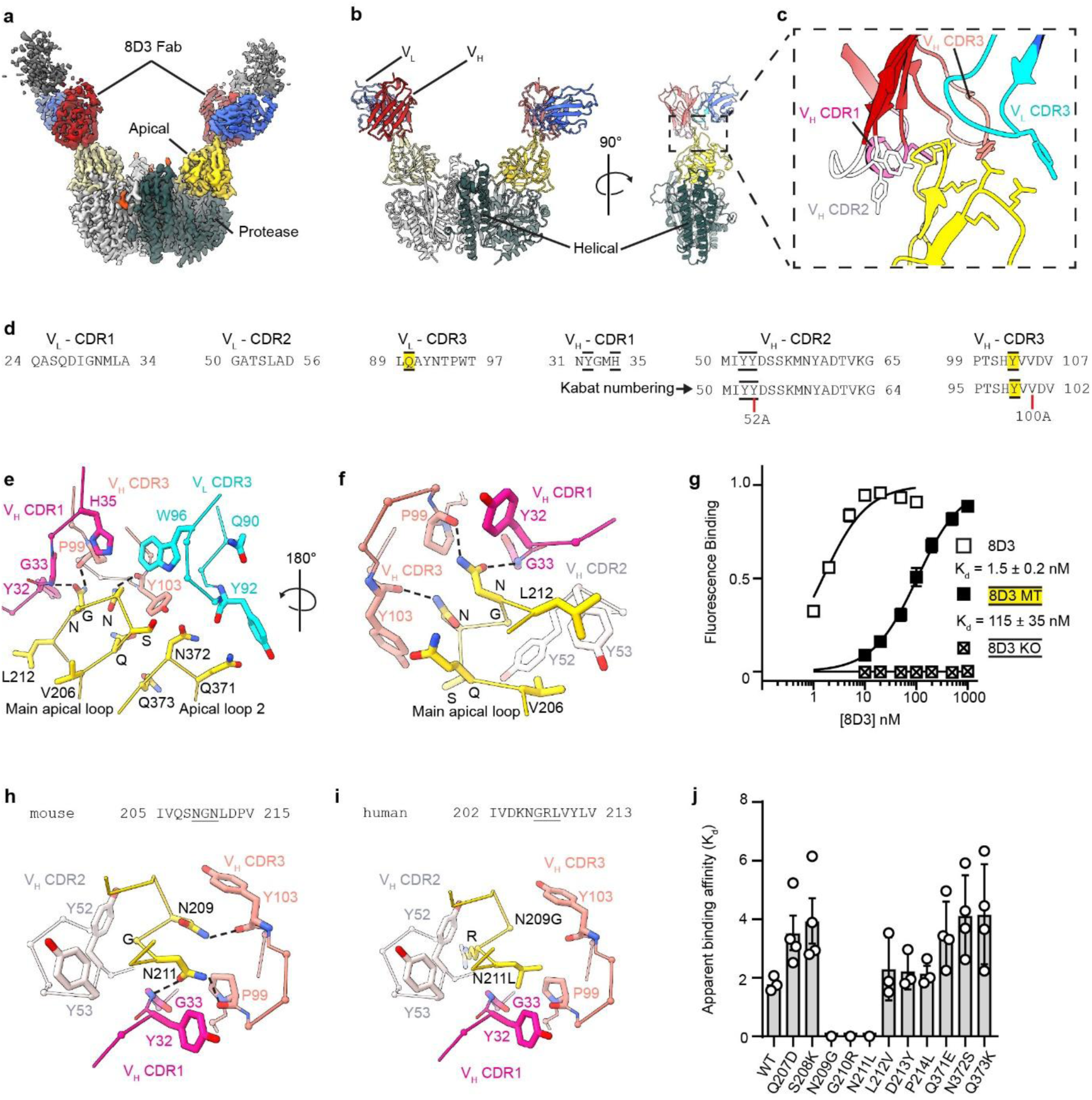
mTfR1 ECD structure and 8D3 binding mode. **a**, Cryo-EM map of mTfR1ECD-8D3 dimeric complex. mTfR1 Protease+helical domain, dark/light grey; mTfR1 apical domain, yellow/pale yellow; 8D3 Fab V_H_ domain, blue; 8D3 Fab V_L_ domain, red; 8D3 Fab C_H_1/C_L_ dark grey; glycans, orange. **b**, Ribbon representation of mTfR1-8D3 dimeric complex structural model. Color scheme same as in **a**. **c**, Close-up of mTfR1-8D3 interacting loops. **d**, Protein sequence of 8D3 Fab CDR loops and Kabat numbering where different for reference. **e**, C_α_-backbone representation of mTfR1-8D3 interacting loops showing key atomic interactions. V_L_ CDR3, cyan; V_H_ CDR1, magenta; V_H_ CDR3, salmon; nitrogen, blue; oxygen, red; dashed lines, putative polar/H-bond interactions (<3.5 Å). **f**, Same as **e**, but rotated 180°. V_H_ CDR2, white. **g**, Fluorescence-based on-cell binding assay measuring 8D3, 8D3-MT (V_L_ Q90A; V_H_ Y103A), and 8D3-KO (V_L_ Q90A/Y92A; V_H_ Y32A/Y52A/Y53A/Y103A) binding to mTfR1-expressing HEK293S stable cells at 4 °C. Apparent binding affinities (*K_d_*; mean ± SEM) determined from *n* = 3 independent experiments. **h**, Close-up of mTfR1 main apical loop N209-G210-N211 interactions with 8D3. Dashed lines, putative polar/H-bond interactions (<3.5 Å). **i**, Same as (G) but NGN replaced with human TfR1 GRL which loses interactions. **j**, Apparent binding affinities of 8D3 for mTfR1 with apical loop mutations. Values are mean ± SEM determined from binding curves from *n* = 3-4 independent experiments.

The mTfR1 architecture is highly similar to hTfR1 (RMSD C_α_ = 0.92Å versus hTfR1 PDB:1CX8), along with high sequence identity, 77 %, and sequence similarity, 88%. However, 8D3 is mouse species specific^43^ and protein sequence alignments reveal low conservation in the main apical binding loop (Extended Data Fig.2b-e). In particular, the mTfR1 N209-G210-N211 binding motif is G207-R208-L209 in hTfR1, losing key contacts from N209 and N211 with the 8D3 peptide backbone, and introducing a potential steric clash from the arginine with V_H_ CDR2 (Fig. 1h and 1i). Consistent with the structural prediction, single substitutions in mTfR1 to the human equivalent for the main loop 207-214 and minor loop 371-373 reduced 8D3 apparent affinities less than 3-fold, except for N209G, G210R or N211L, which all lost binding signal (Fig. 1j). Loss of binding was not due to reduced or ablated mTfR1 surface expression for N209G or N211L, which remained comparable to wild-type (Extended Data Fig. 2f), although G210R reduced surface expression by ∼95 % and thus its direct impact on binding cannot be determined. Nevertheless, single substitutions N209G or N211L both ablate binding to explain mouse versus human species specificity.

### 8D3 is structurally compatible with mTfR1 cross-linking without disrupting receptor distribution

The 8D3 binding mode positions the two Fabs on the mTfR1 dimer pointing away from each other such that the Fab C_H_1-C_L_ Cys-bridges are ∼12.7 nm apart and cannot belong to a single human IgG1 which requires a distance <2.1 nm, ruling out intramolecular avidity where a single antibody double binds a single mTfR1 (Fig. 2a,b). Nevertheless, previous studies report differences of between 5- and 20-fold in apparent affinity between bivalent (BiV) and monovalent (MoV) 8D3 formats, cited to be due to avidity effects ^29,33,55,56^, which can therefore only be explained by intermolecular cross-linking between separate receptors. However, adjacent mTFR1-8D3 complexes place the Fab Cys-bridges at 6.1 nm apart, also precluding intermolecular cross-linking (Fig. 2c). Instead, ∼30° tilting of TfR-ECD domains towards each on the ends of their flexible transmembrane stalks brings the Fab Cys-bridges within range, ∼1.9 nm, without creating any steric clashes between the Fab arms (Fig. 2d). This could permit the intermolecular cross-linking of many receptors together, as proposed for some hTfR1 antibodies which cause clustering, internalisation and degradation ^21,30,34,35^. Alternatively, according to the structural modelling, 8D3 could induce the formation of enclosed paired dimers that would prevent clustering (Fig. 2d). To test for clustering, mouse brain endothelial (bEnd3) cells were incubated for 24h at 37°C with bivalent 8D3 antibody, BiV-8D3, versus monovalent antibody, MoV-8D3, that cannot cross-link or exhibit avidity. Cell surface mTfR1 imaging revealed no difference in the disperse punctate mTfR1 surface distribution by either antibody versus pre-fixed untreated cells (Fig. 2e), showing that any cross-linking if it occurs is limited to small receptor groups, such as pairs, that cannot be detected. Furthermore, incubation with BiV-8D3 versus MoV-8D3 at non-saturating and saturating doses for up to two hours at 37°C in a 96-well plate assay had no impact on quantitative mTfR1 surface expression levels, suggesting no induction of internalisation (Fig. 2f-g). An alternative quantitative assay involving overnight incubation at 37°C and subsequent flow cytometry also revealed no reduction in surface receptor levels across a range of concentrations for both BiV-8D3 and MoV-8D3 versus no antibody treatment (Fig. 2h).

**Fig. 2.**
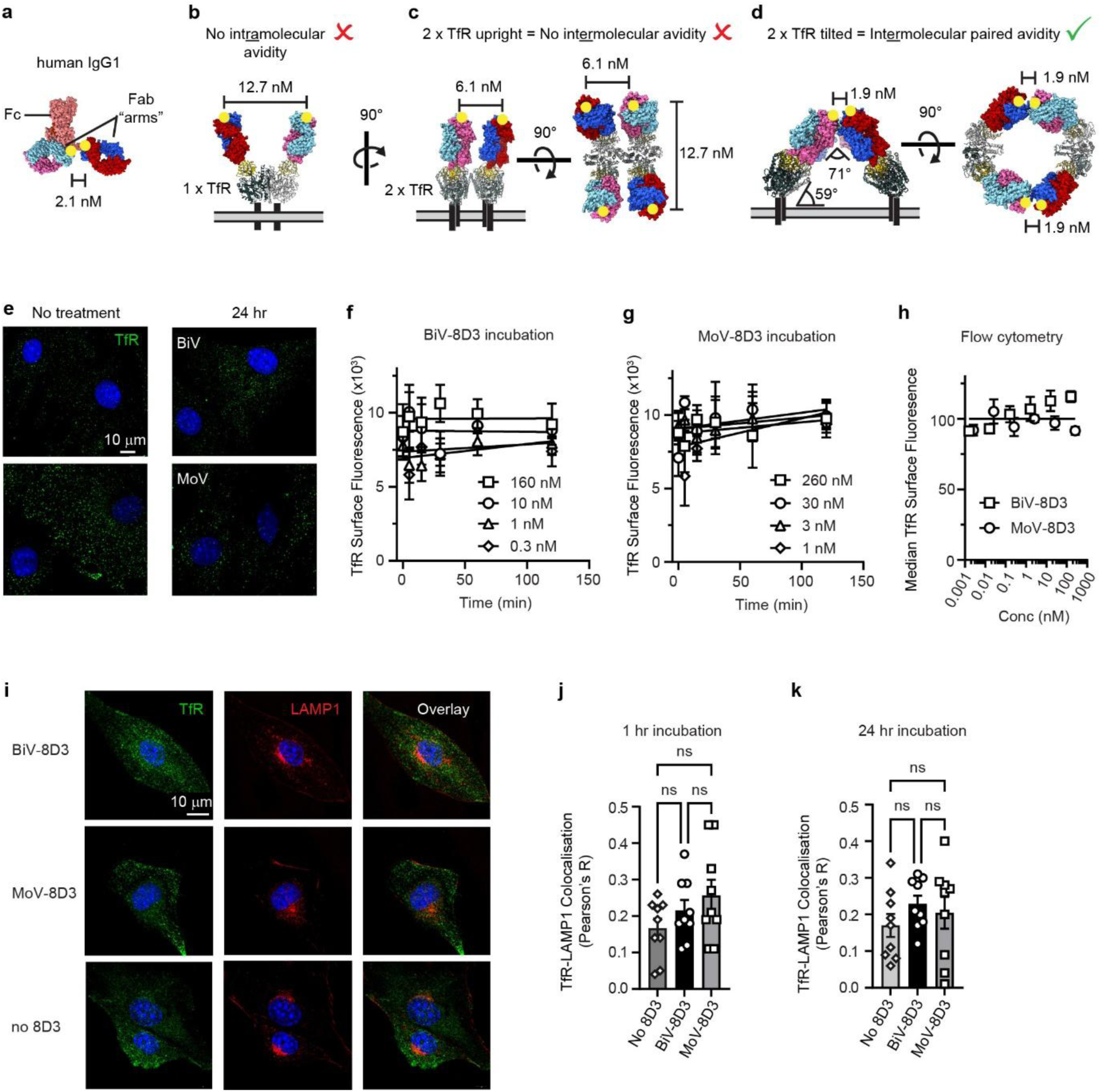
8D3 structural cross-linking mechanism and impact on mTfR1 surface localisation. **a**, Sphere-atom representation of a human IgG1 structure (PDB:1HZH) showing the distance between the (C_H_1-C_L_) Cys-bridges (yellow spots) at the base of each Fab arm. Fab V_H_ domain, blue/light blue; Fab V_L_ domain, red/pink; Fc, salmon. **b**, Structural model showing two 1HZH Fab arms superposed to 8D3(V_L_V_H_)-mTfR1 complex showing the Fab arm Cys-bridge distance, which precludes intramolecular avidity. mTfR1, grey/yellow. **c**, Same as **b**, but showing two adjacent mTfR1. If mTfR1 is upright in the membrane then Fab arm Cys bridges are not in range to come from a single Fc domain regardless of orientation. **d**, Tilting of adjacent mTfR1 towards each other can permit an intermolecular paired avidity. **e**, Representative images of bEnd3 cell-surface mTfR1 staining (green; nuclei, DAPI in blue) by saturating doses of BiV-8D3 or MoV-8D3 either for 24 h at 37 °C prior to fixation or after fixation, showing no obvious mTfR1 clustering at the plasma membrane. From *n* = 3 independent experiments. **f**, and **g**, Time course of bEnd3 surface mTfR1 signal during incubation with varying concentrations of **f**, BiV-8D3, or **g**, MoV-8D3, at 37 °C, showing no detectable reduction in surface mTfR1. Values are mean ± SEM determined from *n* = 5 independent experiment repeats. **h**, Flow cytometry quantification of bEnd3 surface mTfR1 after overnight incubation with varying concentrations of BiV-8D3 or MoV-8D3 at 37 °C. Values are mean ± SEM determined from *n* = 3 independent experiment repeats. **i**, Representative images of mTfR1 (green) and the lysosomal marker LAMP1 (red) in bEnd3 cells incubated with BiV-8D3, MoV-8D3, or medium only (No 8D3) at 37 °C for 24 h. From *n* = 3 independent experiments. **j**, and **k**, Quantification of mTfR1–LAMP1 colocalization expressed as Pearson’s correlation coefficient following incubation with BiV-8D3, MoV-8D3, or medium (No 8D3) at 37°C for 1 h, in **j**, or 24 h, in **k**. Values are mean ± SEM determined from *n* = 9 individual cells from 3 independent experiments; ns, not significant, two-tailed unpaired t-test.

Importantly, the steady state mTfR1 surface expression is not due to an absence of endocytosis in the bEND3 cells because 24-hour incubation at 37°C with BiV-8D3 or MoV-8D3 resulted in fluorescence intensity ∼6-fold higher in permeabilised cells stained for surface plus intracellular receptors than for non-permeabilised cells that detect only surface receptors, confirming constitutive receptor internalisation (Extended Data Fig. 3a,b). In contrast, 8D3-KO treatment was ∼6-fold fainter showing that BiV-8D3 and MoV-8D3 internalisation was not passive and required active binding and uptake with mTfR1 (Extended Data Fig. 3a,b). In addition, intracellular distribution of mTfR1 was comparable between BiV-8D3, MoV-8D3 and 8D3-KO treatments, being dispersed and speckled (Fig. 2i, Extended Data Fig. 3c). BiV-8D3 and MoV-8D3 were highly colocalised with mTfR1 (Pearson’s coefficient ∼ 0.65-0.7), showing they were actively retained to mTfR1 when inside the cell, versus 8D3-KO (Pearson’s coefficient ∼ 0.25) (Extended Data Fig. 3d). They were also more strongly colocalised with the early endosome EEA1 marker (Pearson’s coefficient ∼ 0.35) versus 8D3-KO (Pearson’s coefficient ∼ 0.1) (Extended Data Fig. 3e), with the remainder presumably being bound to mTfR1 at the cell surface or as it passed through other non-early endosomal compartments of the transcytosis machinery. Finally, incubation with saturating doses of BiV-8D3 versus MoV-8D3 for 1 or 24-hours at 37°C did not change mTfR1 colocalization with the lysosomal marker LAMP1 compared with no-8D3 treatment control (Fig. 2i-2k). This suggests that any potential intermolecular cross-linking does not induce measurable lysosomal trafficking or lysosome-associated degradation of surface mTfR1 and contrasts with observations for alternative TfR1 antibodies^21,30,34,35^. Of note, a previous study did report that incubation with BiV-8D3 reduced mTfR1 surface levels^12^. However, we show here that the detection antibody used in that study (RI7) competes with 8D3 for binding to mTfR1 (Extended Data Fig. 3f,g). Thus, the reduction in RI7 staining of surface receptors previously reported can instead be explained by direct blockade of RI7 binding to mTfR1 by pre-bound 8D3 rather than 8D3 reducing mTfR1 surface levels.

### Despite normal mTfR1 distribution 8D3 exhibits inter-receptor avidity in bEND3 cells

Despite an absence of effect by BiV-8D3 on mTfR1 surface distribution, internalisation or degradation, a mouse bEnd3 on-cell binding assay performed at 37°C did reveal a significant 10-fold increased apparent affinity for mTfR1 by BiV-8D3 versus MoV-8D3 (Fig. 3a), consistent with a previous on-cell binding report^56^, and suggesting that there is an intermolecular cross-linking avidity effect. An alternative assay format to measure dissociation from bEnd3 cells at 37°C also revealed an ∼6-fold reduced dissociation rate for BiV-8D3, again consistent with an intermolecular cross-linking avidity effect (Fig. 3b). Under the scenario that BiV-8D3 molecules participate in intermolecular avidity then each antibody can occupy two mTfR1 ECDs versus only one for MoV-8D3, and so the total number of Fc domains bound to cells will be lower for BiV-8D3 (Fig. 3c), although it will not expected to be fully reduced to half the amount under saturation binding conditions because BiV-8D3 molecules will compete for single versus double occupancy. Consistent with this, quantification of the total surface bound Fc domains by flow cytometry using an anti-Fc antibody revealed a significantly lower total binding for BiV-8D3 (Fig. 3d). To further investigate avidity, we performed these assays on a different cell line, HEK cells stably expressing mTfR1, which showed a 10-fold higher density of surface mTfR1 (Fig. 3e, Extended Data Fig. 4a,b). In contrast to the bEnd3 cells, there was only a 3-fold increase in apparent affinity (Fig. 3f), and no significant difference in dissociation rate (Fig. 3g) or total antibody binding (Fig. 3h), revealing that the avidity effect was not obviously detectable in this cell line. To explore why inter-receptor avidity differs between cell lines, we measured mTfR1 colocalization with clathrin heavy chain (CHC) in both cell types. mTfR1 transcytosis is dependent on localisation to clathrin coated pits (CCPs), where each receptor is spatially separated at anchor points by direct interactions with clathrin associated protein AP-2 molecules within the clathrin lattice ^57^. bEnd3 cells exhibited low total CHC expression with no obvious surface delineation (Fig. 3i), in agreement with previous reports that the brain endothelium limits CCP transcytosis to maintain brain barrier integrity^7,58^. In contrast, the HEK293 cell line exhibited strong CHC staining that demarcated the cell membrane boundary (Fig. 3i) and showed a significantly higher colocalization between mTfR1 and CHC (Fig. 3j). Thus, in cells where mTfR1 is anchored at CCPs, intermolecular cross-linking and avidity by BiV-8D3 are reduced (Fig. 3k). However, in bEnd3 cells which have low CCP expression and mTfR1 anchoring, the mTfR1environment is more permissive for mTfR1 cross-linking and avidity.

**Fig. 3.**
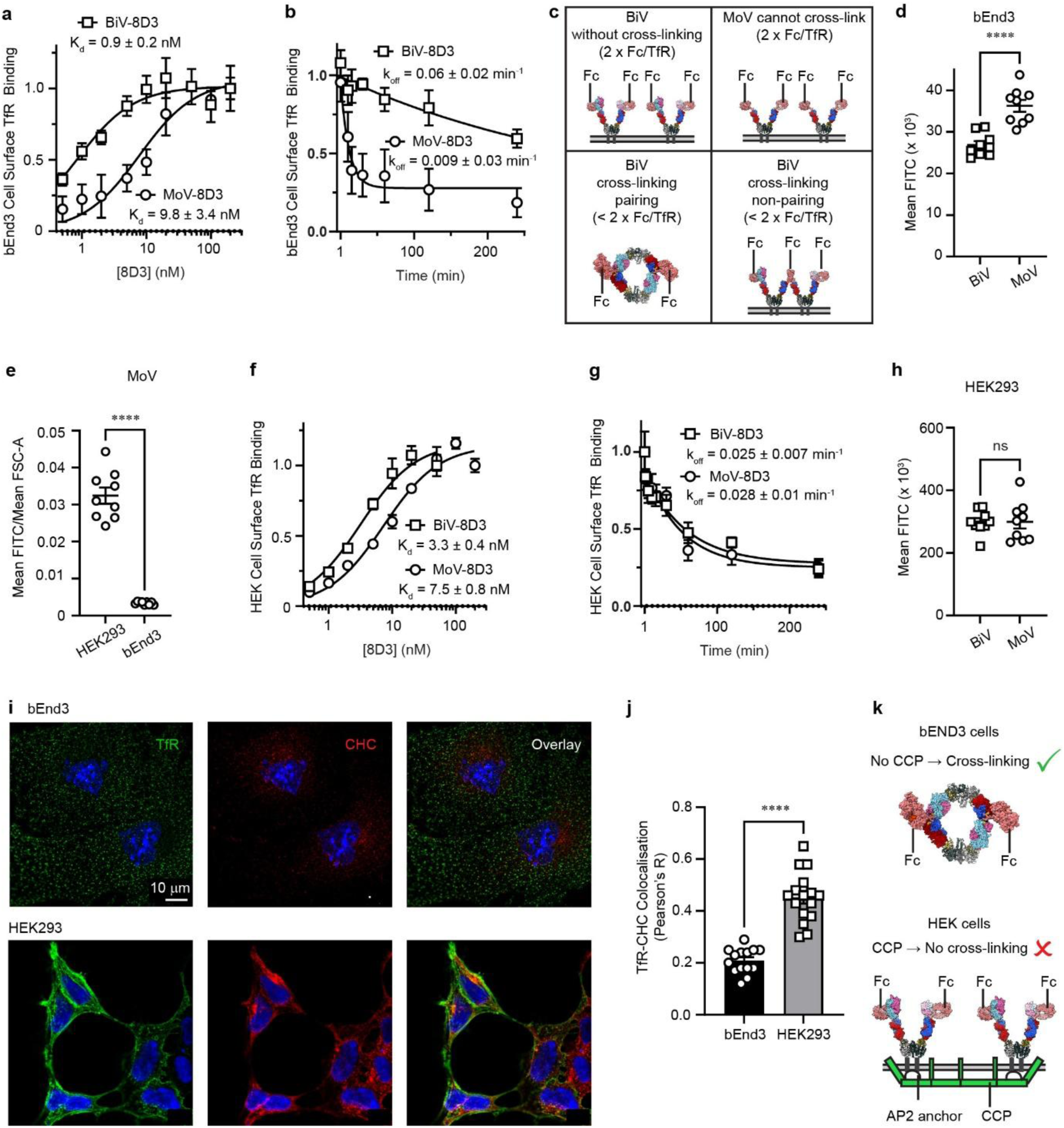
Binding properties of 8D3 for cell surface mTfR1. **a**, Fluorescence-based on-cell binding assay for bEnd3 cells at 37 °C. *K_d_* (mean ± SEM) determined from binding curves from *n* = 4 independent experiments. **b**, Dissociation kinetics from bEnd3 cells at 37 °C. Dissociation constant, *k_off_* (mean ± SEM) determined from *n* = 5 independent experiments. **c**, Structural models of Fc domain binding load on mTfR1 at the cell surface. Without cross-linking, four copies of BiV-8D3/MoV-8D3 bind two receptors in the absence of cross-linking giving four Fc domain staining labels, but only two bind for paired cross-linking, and three for non-paired cross-linking, both of which predict reduced staining intensity for BiV staining, as observed in, **d**. **d**, Corresponding flow cytometry quantification of total antibody bound to bEnd3 cell-surface (detected with DyLight488 anti-human Fc) showing reduced BiV-8D3 versus MoV-8D3 staining (mean ± SEM; BiV-8D3, 26.86 ± 0.92; MoV-8D3, 36.29 ± 1.43; ****, *P* < 0.0001, two-tailed unpaired t-test, *n* = 9 repeats from 3 independent experiments). **e**, Flow cytometry total surface mTfR1 expression (fluorescence normalised to mean FSC-A to account for cell size). Values are mean ± SEM; HEK293, 0.032 ± 0.0022; bEnd3, 0.0034 ± 0.00014; ****, *P* < 0.0001, two-tailed unpaired t-test, *n* = 9 repeats from 3 independent experiments. **f**, and **g**, Same as **a**, and **b**, respectively but for HEK293 cells stable expressing mTfR1 at 37 °C. Values are mean ± SEM from *n* = 6 independent experiments. **h**, Same as, **d**, but for HEK293 cells stable expressing mTfR1. Mean ± SEM; BiV-8D3, 299 ± 12.5; MoV-8D3, 299 ± 21.2; ns, not significant, two-tailed unpaired t-test, from *n* = 9 repeats from 3 independent experiments. **i**, Representative images of mTfR1 (green) and clathrin heavy chain (CHC; red) in bEnd3 and HEK293 cells, from *n* = 3 independent experiments. **j**, Quantification of mTfR1–CHC colocalisation from (I), reported as Pearson’s correlation coefficient (mean ± SEM; bEnd3, 0.21 ± 0.013, *n* = 14; HEK293, 0.45 ± 0.024, *n* = 15 individual cells from *n* = 3 independent experiments; ****, *P* < 0.0001, two-tailed unpaired t-test). **k**, Schematic showing that mTfR1 anchored via AP-2 in CCPs in HEK cells cannot be cross-linked by BiV-8D3, contrasting with the scenario in bEND3 cells where mTfR1 is less colocalised with CCPs.

### 8D3 combinatorial histidine scanning reveals graded pH selective binding

Previous studies have shown that histidine protonation can induce pH-sensitive binding between physiological pH7.4 and endosomal pH5.5^59^, so we performed structure-guided histidine scanning mutagenesis of the 8D3 CDRs to develop proof-of-concept antibodies to investigate pH sensitive brain delivery. In addition to a standard approach comparing antibodies each with a single histidine mutation at a different position throughout the antibody CDRs, we applied a combinatorial approach of co-expressing pairs of histidine mutations, one each from the V_L_ and V_H_ chains to increase the number of permutations explored. However, we used the structure to focus this approach to only the four CDR loops directly contacting mTfR1, V_L_ CDR3 with V_H_ CDRs 1-3. This reduced the search space from ∼450 combinations to ∼230, making it more experimentally feasible to apply. All 230 variants were genetically fused to a C-terminal mNeonGreen (mNG) fluorescent tag for rapid approximation of secretion levels in the Expi293F media, yielding mean concentration ∼500 nM, and SDS-PAGE spot checking confirmed correct band sizes and an absence of protein cleavage/degradation products (Extended Data Fig. 4a-b). Serial dilutions of the raw antibody media for all 230 variants were screened at pH5.5 and 7.4 against the stable mTfR1 HEK293 cell line at 4 °C in 384-well plate format, to generate binding curves. At physiological pH, single histidine mutations across the 9 V_L_ CDR3 positions and 22 V_H_ CDR 1-3 positions were well-tolerated versus wild-type, with no substitutions increasing sensitivity relative to wild-type, and none reducing sensitivity by >3-fold (Fig. 4a and 4b). At physiological pH, double histidine mutations were also well-tolerated, generally remaining within 2-fold of wild-type with the notable exception of V_L_ CDR3 W96H and V_H_ CDR3 P99H, which when combined with other His mutations gave multiple instances of reduced sensitivity >5-fold (6/22 for W96H and 8/9 for V_H_ CDR3 P99H)(Fig. 4c and 4d). Manual inspections of curve fit profiles at physiological pH7.4 versus pH5.5 (chosen to approximate endosomal pH) revealed that most single and double substitution profiles were indistinguishable (Extended Data Fig. 5c,d). For instances where there were differences, these were most clearly manifest as a change in the maximal binding signal rather than the curve fit (∼K_d_) so this metric was chosen for analysis, applying an arbitrary 20% cut-off in difference as being meaningful. For V_L_ CDR3 mutations, none alone (co-expressed with V_H_ WT) caused a pH-dependent change in maximal binding (Fig. 4e). However, for V_H_, although most mutations alone (co-expressed with V_L_ WT) had no effect, CDR2 Y53H or CDR3 P99H, did increase binding at pH5.5 by 36% and 93% respectively (Fig. 4f,g). For the double His mutant combinations, various combinations increased maximal binding at pH5.5 even though the equivalent single mutations alone had had no effect (Extended Data Fig. 5f,g). For V_L_ CDR3, W96H most frequently combined with V_H_ mutations (7/22) to favour pH5.5 binding (Fig. 4f,h, Extended Data Fig. 5f). For V_H_, the substitutions that most frequently combined with V_L_ CDR3 substitutions to favour pH5.5 binding were V_H_ CDR3 Y103H (5/9) and D106H (3/9)(Extended Data Fig. 5g). In the case of V_H_ CDR3 P99H, the only substitution to have an effect alone, when expressed in combination with V_L_ substitutions it consistently produced 40-400% increases in maximal binding at pH5.5 (Fig. 4e). Importantly, in all cases pH preference favoured increased maximal binding at pH5.5, none favoured binding at pH7.4.

**Fig. 4.**
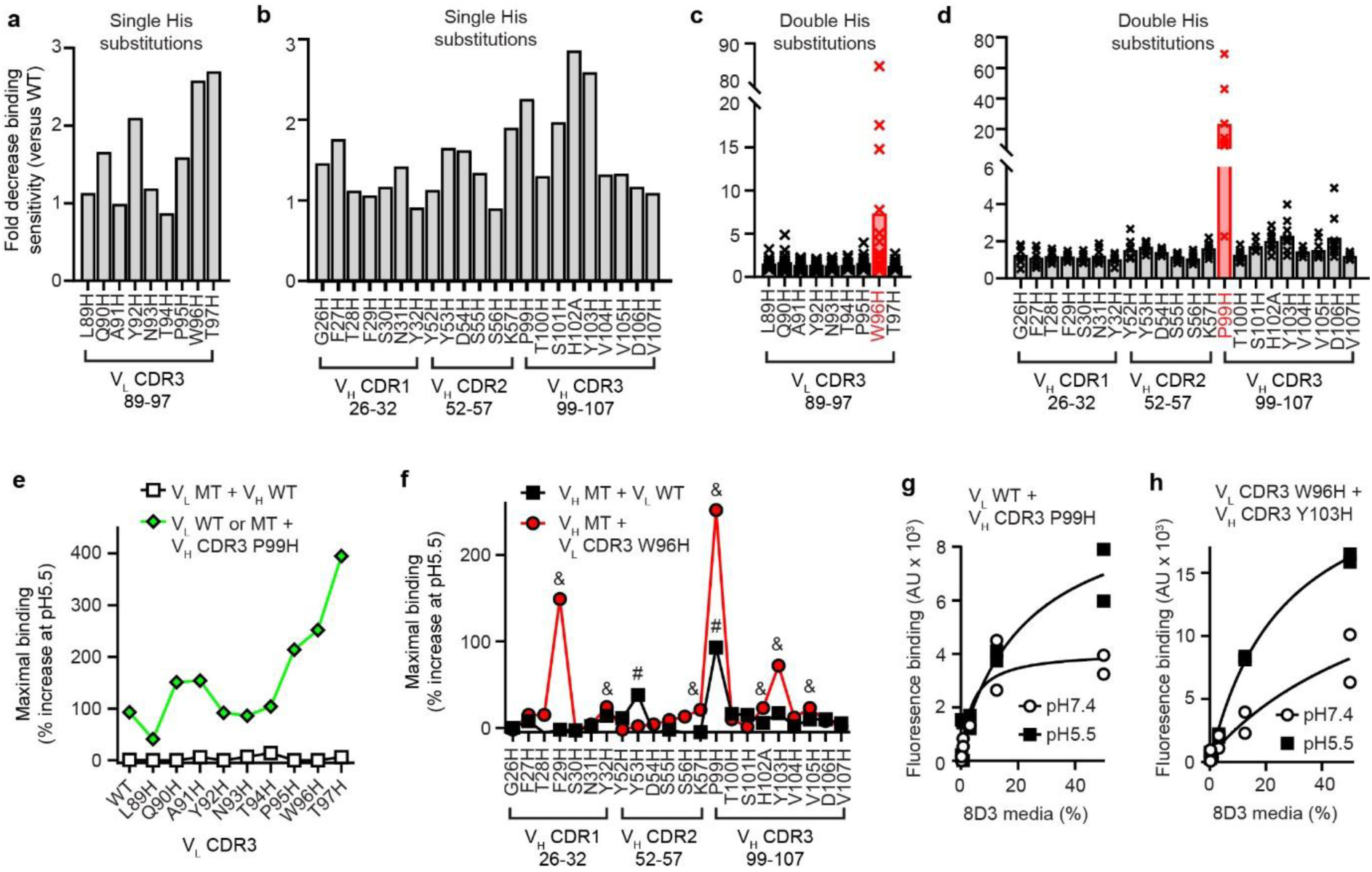
pH-sensitive 8D3 library screening. **a**, and **b**, Cell-surface mTfR1 binding sensitivity at pH 7.4 for 8D3 variants with single histidine substitutions in **a**, V_L_, or **b**, V_H_. **c**, and **d**, Cell-surface mTfR1 binding sensitivity at pH 7.4 for 8D3 variants with double histidine substitutions, one in each V_L_ and V_H_. **c**, V_L_ substitutions show a data point for each paired V_H_ substitution. **d**, V_H_ substitutions show a data point for each paired V_L_ substitution. Red bars indicate the V_L_ or V_H_ substitution that reduces binding sensitivity the most upon pairing. **e**, Increase in maximal 8D3 binding at pH5.5 for single V_L_ substitutions versus in combination with V_H_ P99H. **f**, Increase in maximal 8D3 binding at pH5.5 for single V_H_ substitutions versus in combination with V_L_ W96H. # >20% maximal binding at pH5.5 for single V_H_ substitutions; & >20% maximal binding at pH5.5 for V_L_ W96H + V_H_ substitutions. **g**, Representative binding curves at pH5.5 versus pH7.4 for 8D3 with V_H_ P99H mutation. **h**, Representative binding curves at pH5.5 versus pH7.4 for 8D3 with V_L_ W96H V_H_ P99H mutations. In all experiments screening was done as *n* = 1 experiment (technical duplicates).

We selected several variants that represented a range of strengths of effect to express and purify to obtain apparent K_d_s, in order to produce a panel of 8D3 antibodies with progressively graded sensitivities to pH-dependent binding. As previously observed from the crude media screening, for the purified proteins single substitutions in residues V_L_ W96A, V_H_ Y52H, Y103H and D106H also did not reduce apparent affinity by more than 2-fold or exhibit pH sensitive binding (Fig. 5a, Supplementary Table 2), and V_H_ P99H showed pH5.5 preferential binding (Fig. 5b,c, Supplementary Table 2). Furthermore, all four double substitutions engendered pH5.5 preferential binding spanning a range of apparent K_d_s. V_L_ W96H V_H_ D106H retained WT apparent affinity at pH5.5 (K_d_ 2.3 nM), which was reduced 4-fold at pH7.4 (K_d_ 8.3 nM) (Fig. 5a, Extended Data Fig. 6a; Supplementary Table 2). For V_L_ W96H V_H_ Y52H the pH5.5 K_d_ versus WT increased 8-fold to 12 nM and achieved 5-fold selectivity versus pH7.4 (K_d_ 65 nM)(Fig. 5b, Extended Data Fig. 6b; Supplementary Table 2). For V_L_ W96H V_H_ Y103H the pH5.5 K_d_ increased 5-fold versus WT to 7.0 nM, with 20-fold selectivity versus pH7.4 (K_d_ 142 nm), and for V_L_ W96H V_H_ P99H the pH5.5 K_d_ increased 9-fold to 15 nM, and no binding was detectable at pH7.4, resulting in >> 13-fold selectivity (Fig. 5B and S6C and S6D; Table S2).

**Fig. 5.**
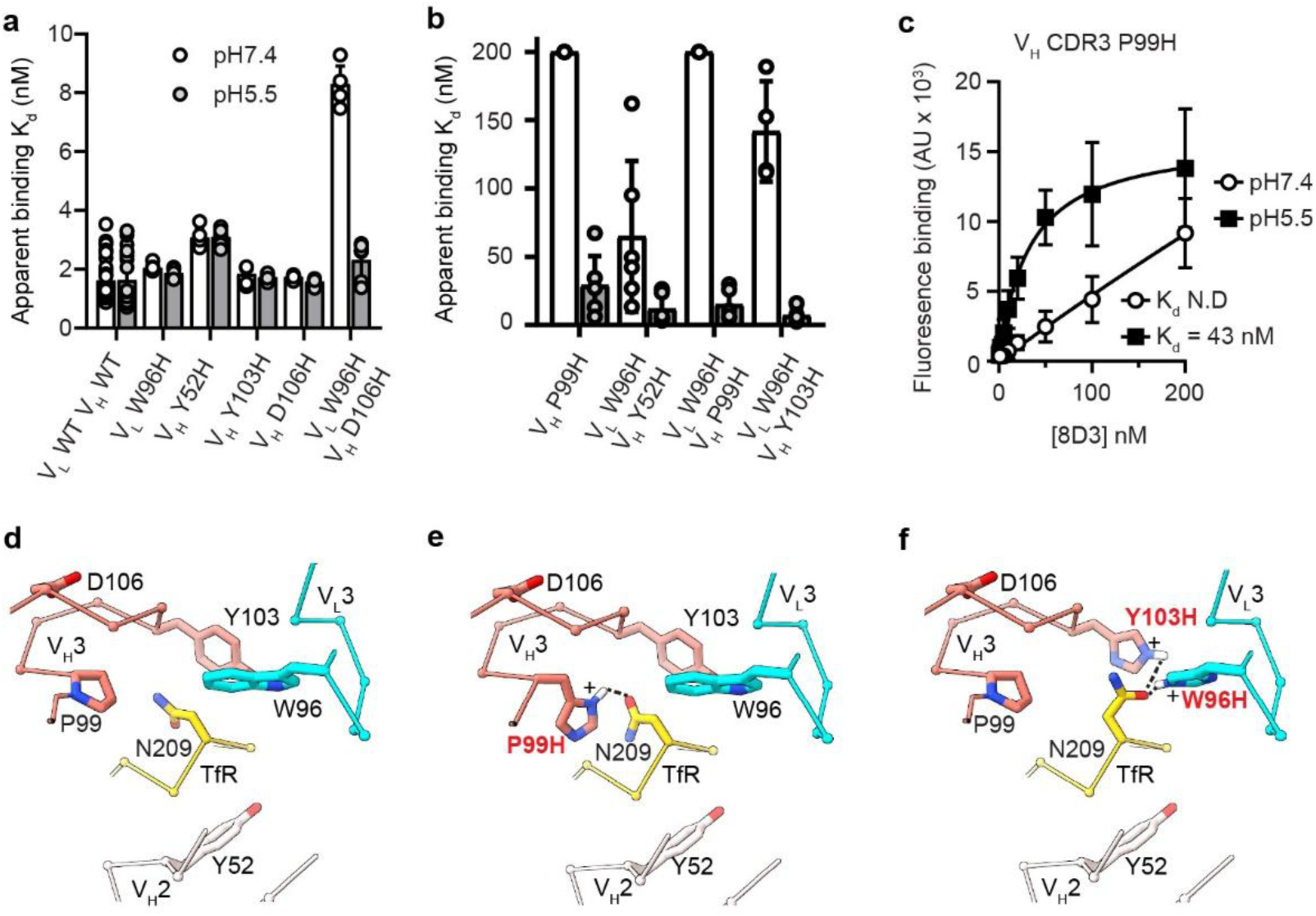
Purified binder pH-sensitivity and binding mechanism modelling. **a**, and **b**, Apparent binding affinities (*K_d_*; mean ± SEM) determined using purified 8D3 variants with indicated histidine substitutions, measured on mTfR1-expressing HEK293 cells at pH 7.4 and pH 5.5, from *n* = 28 for wild-type, *n* = 6 for mutations (except 8D3 V_L_ W96H V_H_ Y103H pH7.4, *n* = 4), independent experiments. **b**, is for mutations that caused larger shifts in pH sensitivity. **c**, On-cell binding curves at pH5.5 versus pH7.4 for the 8D3 V_H_ P99H variant. Values are mean ± SEM determined from binding curves from *n* = 6 independent experiments. **d-f**, C_α_ and side chain stick representations of mTfR1 main apical loop at N209 and 8D3 binding loops V_L_3, V_H_2 and V_H_3 for **d**, wild-type, versus **e**, 8D3 V_H_ P99H and **f**, 8D3 V_L_ W96H V_H_ Y103H. Nitrogen, blue, oxygen, red, hydrogen, white. For each modelled histidine mutation position (common rotamers used), protonation (+) at pH5.5 is positioned to be able to interact with N209 to influence pH-dependent binding.

A structure of mTfR1 at pH5.5 bound by an 8D3 His variant could not be solved due to mTfR1 precipitation in low pH buffer. Nevertheless, the structure at pH7.4 bound by WT 8D3 reveals that for each pH sensitive binder there is at least one histidine facing mTfR1 N209, coming either from V_H_ CDR3 P99H or Y103H, or from V_L_ W96H (Fig. 5d-f). Given that N209 is a key binding residue to wild-type 8D3 (Fig. 1j), histidine mutations such as at P99H, or W96H/Y103H (Fig. 5e,f) were they to interact with N209 could destabilize binding at pH7.4 more than at pH5.5 to engender low pH preferential binding. The structure provides a common molecular mechanism to explain this whereby protonation of histidine at pH5.5 converts a neutral hydrogen donor into a positively charged one resulting in donation of a charge-assisted hydrogen bond to the asparagine electronegative carbonyl group, strengthening the interaction (Fig. 5e,f). In the case of the more distant residues V_H_ CDR2 Y52H or V_H_ CDR3 D106, these can act indirectly on N209 through interactions with the back of the N209 loop or the back of the V_H_ CDR3 loop respectively (Fig. 5d), which alone have no effect but in combination with W96H, do cause modest reductions in affinity which are largely recovered at low pH (Fig. 5a,b).

### Preferential TfR1-8D3 interactions at low pH prevent brain penetration of 8D3

Previously, two antibodies reported to preferentially bind hTfR1 at pH7.4 over endosomal pH5.5 showed superior transcytosis in an *in vitro* BBB model^36^. However, this was not tested *in vivo*, and in our pH-dependent binding assay we did not detect pH sensitivity for either antibody (Extended Data Fig. 6e,f). Another study on nanobodies suggested stronger binding at pH7.4 than pH5.5 enhances brain penetration, although the correlation between pH sensitivity and brain penetration was low^60^. Finally, effective brain delivery has been shown for a pH sensitive antibody, however, this effect was attributed to a pH-sensitive conformation of TfR1 ^30^. As a result, antibodies with equivalent binding modes that lack pH sensitivity cannot be developed and compared to quantify the contribution of pH-dependent binding to brain delivery for that antibody. To confirm if pH preferential antibody binding does effect brain penetration, we collected *in vivo* data for the pH-sensitive 8D3 antibodies. We performed ELISA-based quantification of the antibodies from mouse plasma and brain parenchyma 24 h after intraperitoneal (IP) injection. As with all antibodies used throughout this study, Fc effector-function was absent to minimise immune-mediated effects^61^. Plasma concentrations of both high affinity binders, BiV-8D3 and MoV-8D3, were significantly lower than the negative control, BiV-8D3-KO (Fig. 6a, Supplementary Table 3). This is consistent with target-mediated drug disposition, often reported for high-affinity TfR1 shuttles due to uptake via peripheral TfR1 increasing plasma clearance^30,61^. In contrast, the low affinity binder, BiV-8D3-MT, and the three pH sensitive 8D3s, pH1 (V_L_ W96H V_H_ Y52H), pH2 (V_L_ WT V_H_ P99H), and pH3 (V_L_ W96H V_H_ P99H) were not significantly reduced versus BiV-8D3-KO (Fig. 6a, Supplementary Table 3). Regardless of any disposition effects, plasma concentrations for BiV-8D3, MoV-8D3, BiV-8D3-MT and BiV-8D3-pH1, were all sufficiently high to reach predicted occupancies for mTfR1 on the BBB luminal side of ∼50 % or greater (Fig. 6b), to ensure engagement for transcytosis into the brain. Despite a predicted ∼100 % mTfR1 engagement from the luminal side, BiV-8D3 only showed a non-significant trend towards a higher brain concentration than BiV-8D3-KO (Fig. 6c). In contrast, MoV-8D3 and BiV-8D3-MT had lower predicted luminal engagement, ∼70% and ∼50% respectively, due to having lower affinities, but both achieved brain concentrations significantly higher than BiV-8D3-KO (Fig. 6c). This is consistent with previous studies showing that reducing valency or affinity of bivalent antibodies can both improve release and so delivery into the brain parenchyma ^21–23,26^. These brain concentrations corresponded to predicted occupancies for mTfR1 in the brain (on the parenchymal side of the BBB) by BiV-8D3 and MoV-8D3 of ∼70% and ∼ 60% respectively, substantially higher than for BiV-8D3-MT, 3% (Fig. 6d). For the three pH sensitive antibodies, parenchymal concentrations were all similar to BiV-8D3-KO, being less than 1 nM, and all were significantly lower than the MoV-8D3 and BiV-8D3-MT (Fig. 6d). Most notably, BiV-8D3-pH1 has apparent K_d_ 65 nM at pH7.4, and luminal mTfR1 occupancy of 62 %, both values in between those of the BiV-8D3/MoV-8D3 and BiV-8D3-MT, meaning it would ordinarily be expected be elevated in the parenchyma, showed no sign of mTfR1-dependent brain delivery, suggesting that preferential low pH binding prevents brain delivery (Fig. 6c).

**Fig. 6.**
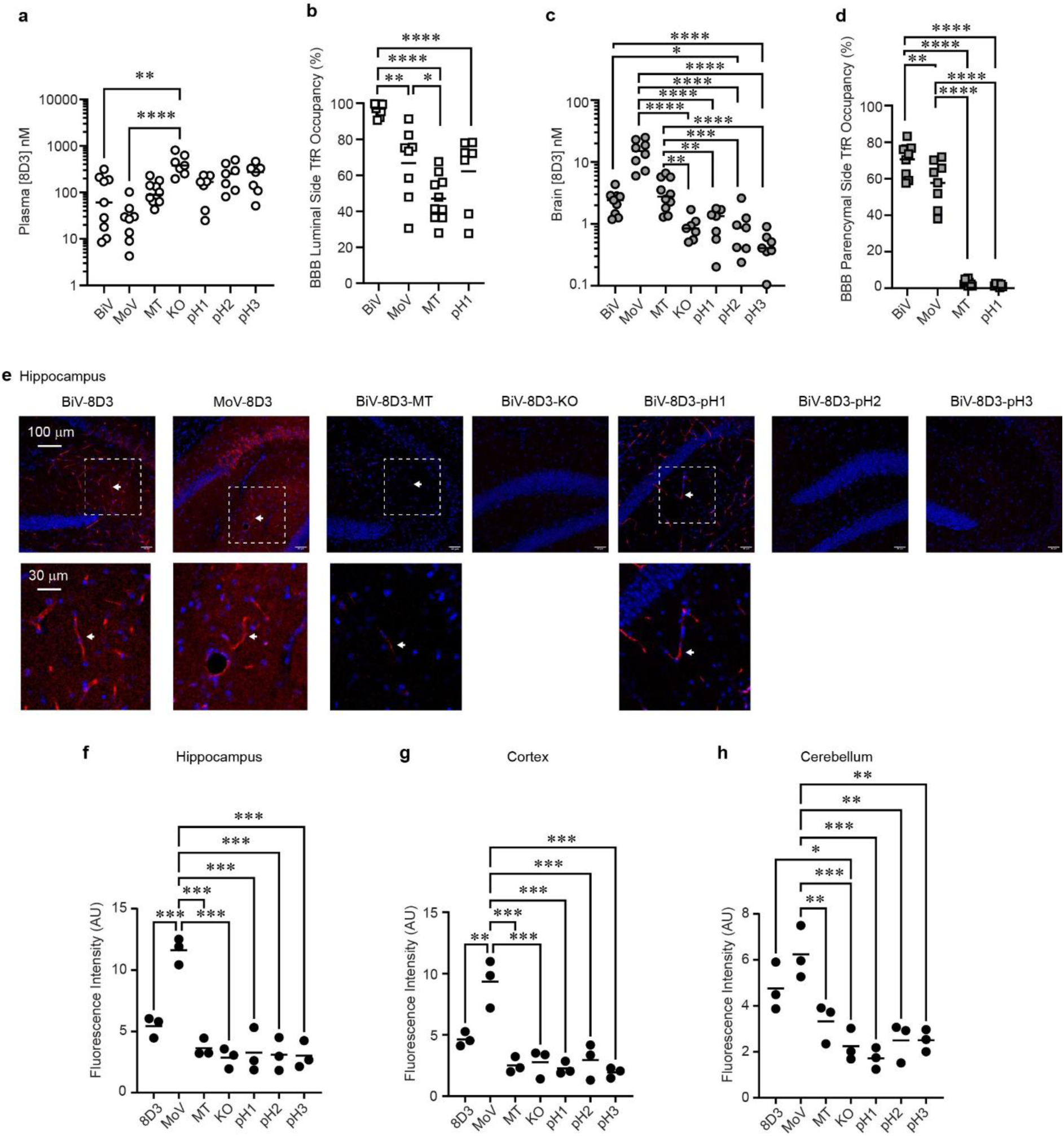
*in vivo* distribution and brain penetration. **a-d**, ELISA quantification of 8D3 antibody concentrations in **a**, plasma, and **c**, brain, and corresponding mTfR1 receptor occupancy at **b**, the BBB luminal side, and **d**, abluminal/parenchymal side. Occupancies were calculated from plasma/brain concentrations versus corresponding apparent affinities. BiV-8D3, *n* = 9; MoV-8D3, *n* = 8; 8D3-MT, *n* = 10; 8D3-KO, *n* = 7; 8D3-pH1-3, *n* = 7, mice per group. **e**, Representative sagittal brain sections showing distribution of 8D3 antibodies in hippocampus at 24 h after intraperitoneal injection (20 mg/kg). 8D3, red (anti-human IgG staining); nuclei, blue (DAPI). Dashed boxes indicate regions enlarged below; arrows mark vessel staining. **f-h**, Quantification of parenchymal 8D3 signal from confocal images of **f**, hippocampus, **g**, cortex, and **h**, cerebellum. Fluorescence intensity (AU) per section was measured from five randomly selected 10 × 10 pixel ROIs per image placed in parenchyma while avoiding vessel-associated staining (*n* = 3 mice per group). Values are mean ± SEM. Statistical comparisons: one-way ANOVA with Tukey’s multiple-comparisons test; *, *P* < 0.05; **, *P* < 0.01; ***, *P* < 0.001; ****, *P* < 0.0001.

To complement the ELISA data we confirmed direct antibody engagement of BBB mTfR1 and release into the parenchyma by immunohistochemistry. This revealed that BiV-8D3, MoV-8D3, BiV-8D3-MT and BiV-8D3-pH1 were bound to the blood vessels throughout the hippocampus, cortex and cerebellum (Fig. 6e, Extended Data Fig 7a,b), as was shown based on the predicted luminal mTfR1 occupancies (Fig. 6b). This confirms that BiV-8D3-pH1 directly engages the BBB even though it is excluded from entering the brain. Notably, the BiV-8D3-MT staining was fainter, consistent with its slightly lower luminal occupancy and its lower affinity causing wash-off prior to fixation of the tissue. 8D3-KO, BiV-8D3-pH2 and BiV-8D3-pH3, which exhibit no or very low binding at pH7.4 could not be observed on blood vessels, except for some faint staining for BiV-8D3-pH3 in the cortex (Fig. 6e, Extended Data Fig. 7a,b). MoV-8D3 showed visible release from the BBB into the brain parenchyma, confirmed by significantly elevated parenchymal fluorescence intensity versus all other 8D3 variants across all three brain areas, with one exception being versus BiV-8D3 in the cerebellum (Fig. 6f-h). BiV-8D3 showed a trend towards elevated fluorescence in the parenchyma, although this only reached significance versus the non-binder in the cerebellum (Fig. 6f-h). BiV-8D3-MT was not detectable above the non-binder signal (Fig. 6f-h), however this result is expected based on its low predicted parenchymal mTfR1 occupancy, of just 3% (Fig. 6d), meaning the antibody cannot be cross-linked to mTfR1 during the *in vivo* tissue fixation process and instead it will be washed away. Similarly, the pH1-3 variants were all undetectable by this method (Fig. 6f-h), again as predicted from low parenchymal mTfR1 occupancies of less than 5% (Fig. 6d). Together, the on-cell binding apparent K_d_s, ELISA concentrations, predicted occupancies, blood vessel staining, and parenchymal immunofluorescence show good agreement, consistent with previous studies, and demonstrate that a low pH preference antibody can directly engage with the BBB but is excluded from the parenchyma.

## Discussion

Although hTfR1 structures have been available for over two decades^32^, a mTfR1 structure has remained unavailable despite reported attempts to determine one^60^. Although the overall mTfR1 architecture and sequence are highly similar to hTfR1, there are clear local structural differences, particularly in the key immunogenic apical loops targeted by 8D3, which explain antibody species specificity. In the future, comparing experimental structures from different species could provide deeper insights into binder specificity, both for shuttle antibodies and other proteins, such as viral proteins, that target TfR1^19,49^. Furthermore, given the current lack of rodent/human cross-reactive shuttles the availability of the mTfR1 structure could aid de novo design of cross-species antibody shuttles. This can facilitate translational research by removing the need for transgenic hTfR1 knock-in animals which may have altered receptor behaviour and require time-consuming crossbreeding with disease models^38–42^.

The consensus in the field is that bivalent antibodies use intermolecular avidity to cross-link multiple TfR1s at the cell surface, triggering clustering, internalisation and lysosomal targeting^21,34,35^. In contrast, for the model BBB shuttle, 8D3, we show that full-length bivalent antibody does not appreciably disrupt mTfR1 surface or intracellular distribution, surface expression, or lysosomal colocalization, for the mouse brain endothelial cell line, bEND3. Nevertheless, we observed an avidity effect, consistent with shifts in on-cell apparent affinity previously reported^29^, which cannot be explained by structurally incompatible intramolecular avidity. To reconcile an intermolecular cross-linking avidity effect with unperturbed receptor cell distribution the cryo-EM structure is compatible with enabling self-contained receptor pairs to form, which traffic normally. This mechanism is also consistent with an increased colocalization of mTfR1 in clathrin coated pits coinciding with a reduction in avidity, as occurred in HEK293 cells, because intermolecular crosslinking is impeded.

Although prevailing wisdom in the field supports monovalent delivery strategies, our study suggests molecular binding modes that limit TfR1 cross-linking can mitigate concerns about avidity-induced TfR redistribution. This aligns with a recent report favouring bispecific formats because they persist longer in the plasma, do not require Fc-heterodimer engineering and are easier to purify^30^. Interestingly, our data also demonstrate that intermolecular avidity is not just antibody-dependent but also cell-dependent. This is an important consideration for biodistribution studies because the risk of off-target TfR lysosomal targeting might be reduced by higher CCP rates in most cell types compared to the BBB^7,58^.

The structure also enabled combinatorial histidine-scanning to generate gradations of pH-selectivity that allowed us to systematically show that preferential binding at endosomal pH5.5 versus physiological pH reduces brain penetration. This supports a long-standing hypothesis, challenging to prove for antibodies^30,36,60^, that preferential antibody binding at physiological pH versus endosomal pH5.5 enhances brain delivery, although we were unable to test this directly because we did not discover antibodies with this polarity of pH preference. Nevertheless, we have generated a mTfR1 antibody toolset that can be used in future studies, for example to selectively target mTfR1 in murine cancer models where pH is known to be reduced^62,63^.

To summarise, we demonstrate the importance of the molecular binding mode in determining bivalent brain shuttle behaviour, influencing CNS penetration, and controlling pH preferential binding. In contrast to previously studied TfR1 antibodies, which presumably exhibit a different binding mode, we show that bivalent 8D3 does not disrupt mTfR1 cellular distribution despite exhibiting avidity. These findings support the idea that rational design against specific epitopes can produce next-generation bivalent cross-species binding antibodies that minimise TfR1 disruption, whilst retaining optimal pharmacokinetics and manufacturability.

## Acknowledgments

We gratefully acknowledge Lee Cooper, Steven Hardwick and Dima Chirgadze for EM grid preparation and data collection under grant Wellcome Trust (206171/Z/17/Z; 202905/Z/16/Z). O.E.K – Astra Zeneca PhD studentship. A.A.W, P.S.M – Rosetrees Trust RC-IaT-2020\100003. C.S - BBSRC doctoral award BB/X010899/1. L.A.P and E.St.J.S. - a joint and equal investment from UKRI and Arthritis UK (MR/W002426/1) as part of the ADVANTAGE visceral pain consortium through the Advanced Pain Discovery Platform initiative. T.M – Cancer Research UK grant DRCRPG-May23/100002 to C.Siebold. S.A.N – BB/Y007816/1. W-N.C – BBSRC BB/M024709/1. V.M – Leverhulme Trust ECF-2023-547 and Isaac Newton Trust. J.I.B – MRC MR/W006650/1. J.E.L, C.I.W – Astra Zeneca PLC.

## Author contributions

E.S.J.S, J.E.L, C.I.W, P.S.M: conceptualisation and supervision. S.L, O.E.K, A.A.W, C.S, L.A.P, T.M, E.A.S, S.A.N, W-N.C, V.M, J.I.B: investigation and formal analysis. S.L, O.E.K: visualisation. S.L, O.E.K, P.S.M: writing – original draft. All authors: writing – review and editing.

## Conflict of interest

The authors J.E.L and C.I.W declare the following competing interest: were employees of Astra Zeneca during this study. All other authors report no conflict of interest.

## Data availability

Data available on request from the authors. For DNA and protein sequence information of 8D3 please see the plasmid sequences under the Addgene depositions as listed below: pHLsec biV-8D3 HC - 254707; pHLsec biV-8D3 LC - 254690; pHLsec biV-8D3 HC KO – 254708; pHLsec biV-8D3 LC KO – 254709; pHLsec MoV-8D3 HC – 254711; pHLsec MoV-8D3 Fc – 254710. Atomic model coordinates for mTfR1ECD-8D3 have been deposited in the Protein Data Bank with accession code 29GF. The cryo-EM density map has been deposited in the Electron Microscopy Data Bank with accession code EMD-57148.

## METHODS

### Constructs

For the mTfR1 ectodomain a synthetic human codon-optimised DNA sequence corresponding to Uniprot Q62351, with boundaries SRLYW…NIDNEF, was subcloned into the pHLsec mammalian expression vector^64^ between Age/Kpn and followed by a C-terminal Avi-Tag (GLNDIFEAQKIEWHE), linker and 6 x His tag (GGSGGSHHHHHH). Full-length mTfR1 was sub-cloned into pHR-CMV-TetO2^65^ between the EcoRI and KpnI sites with an N-terminal kozak sequence, and the plasmid contains a C-terminal 3C cleavage site followed by a TwinstrepII tag (LEVLFQGPGGSGSA-WSHPQFEK-GGGSGGGSGGSA-WSHPQFEK) after the Kpn site. Synthetic human-codon optimised 8D3 LC and HC constructs were subcloned into the pHLsec between the AgeI and XhoI restriction sites. LC constructs began and ended DIQM…RGEC. HC constructs began and ended EVQL…SPGK and the truncated F_C_ (hole) began and ended KSCD…SPGK. The HC and truncated F_C_ contained the Effector-dead mutations L234F, L235E and P331S^66^. Other mutations were introduced by overlap PCR. The bivalent HC and monovalent HC (knob) contained a C-terminal AviTag, linker and 6 x His tag.

### mTfR and 8D3 Fab expression, purification and cryo-EM sample preparation

Protein expression was as previously described^67,68^. 800 mL of HEK293-GnTI^−^ cells (ATCC, CRL-3022), which produce proteins with truncated N-linked glycans, Man_5_GlcNAc_2_ were grown in suspension up to a density of 2 × 10^6^ cells per mL in Protein Expression Media (PEM) (Invitrogen), supplemented with 1% (v/v) L-glutamine (Gibco), 1% (v/v) non-essential amino acids (Gibco) and 1% (v/v) foetal calf serum (Sigma-Aldrich). Cultures were grown shaking at 130 rpm, 37 °C, 8% CO_2_. For transient transfection, the cells were collected by centrifugation (200 x *g*, 5 min) and resuspended in 100 mL Freestyle media (Invitrogen) containing 1.2 mg PEI Max (Polysciences) and 0.4 mg plasmid DNA. The cells were incubated in the transfection mixture for 4 h, with 160 rpm shaking, 37 °C, 8% CO_2_. After this, PEM medium was added up to 800 mL and the cells were returned to 130 rpm shaking incubator. Protein expression was continued for 5 days, after which the protein-containing media was isolated (the cells were pelleted at 200 x *g*, 5 min) and kept in the cold room for further use.

The protein was purified by nickel affinity chromatography followed by size-exclusion chromatography (SEC). Protein-containing media were diluted two-fold with Phosphate-buffer saline (PBS), followed by addition of 10x binding buffer: 500 mM Tris-HCl pH 8, 3 M NaCl and 3 mL Ni-NTA agarose loose resin (Qiagen). The sample was batch-bound for 1 h at 10 °C and then loaded on an empty column (BioRad). The resin was washed with 50 column volumes (CV) PBS containing 10 mM Tris-HCl pH 8 and 0.05% (v/v) Tween 20, followed by 50 CV PBS containing 10 mM Tris-HCl pH 8, 10 mM imidazole pH 8. One 5 mL elution was performed with PBS containing 20 mM Tris-HCl pH 8, 500 mM NaCl, 250 mM imidazole pH 8. The 8D3 Fab elution was concentration dialysed into PBS to 10 mg/mL using a 10 kDa molecular weight cut off concentrator (Amicon). The 8D3 Fab was mixed at a molar ratio of 1:2 into the mTfR elution and the protein was concentrated slowly at 12 °C in a 10 kDa molecular weight cut off concentrator (Amicon) to ∼6 mg/mL. Protein was subjected to SEC on a Superose 6 10/300GL (GE Healthcare) into PBS supplemented with 2 mM Tris-HCl pH 8, 50 mM NaCl, 25 mM imidazole pH 8. The peak fractions corresponding to the ectodomain dimer were supplemented with an extra 0.3 molar ratio of Fab to ensure saturation of the mTfR1 and concentrated with 100 kDa molecular weight cut off concentrator (Vivaspin) to 1.8 mg/mL. Finally, the sample was diluted 1:2 with 33 % PBS containing 10 mM imidazole to give final protein concentration 0.6 mg/mL. For cryo-EM grid preparation, 3uL of sample was applied on glow-discharged gold R 2/2 200 mesh UltraAuFoil grids (Quantifoil) and then blotted for 3s, with blotting force −3, before plunge-freezing in liquid ethane cooled by liquid nitrogen. Plunge-freezing was performed using a Vitrobot Mark IV (Thermo Fisher Scientific) at approximately 100% humidity and 4 °C.

### Cryo-EM data collection

The cryo-EM dataset was collected on a Titan Krios G3 microscope at the Department of Biochemistry EM facility (BiocEM, University of Cambridge). The microscope was equipped with a Gatan K3 camera and Gatan BioQuantum energy filter. The dataset was acquired automatically using EPU software (Thermo Fisher Scientific, version 3.7–3.9). Details of data collection are summarised in Supplementary Table 1.

### Cryo-EM image processing

A general image processing workflow is show in Extended Data Fig. 1. Motion correction, contrast transfer function correction, and particle picking were performed using RELION (version3.1)^69^. All subsequent particle processing was done in cryoSPARC (version 4.7.1)^70^. Particles were initially picked using the CryoSPARC blob picker and subjected to 2D classification. Selected classes were used to generate templates for template-based picking. The resulting particles were extracted and further cleaned through additional rounds of 2D classification, and selected high-quality classes were used to generate three ab initio models. The best-resolved ab-initio 3D model with the most particles was selected and subjected to homogenous refinement followed by non-uniform refinement to generate an initial reference model. In parallel, poorly resolved classes from earlier rounds of 2D classification were used to generate ‘junk’ ab initio models. All extracted particles were included in three or more iterative rounds of heterogenous refinement using the refined receptor model together with junk classes as references. After iterative classification, only particles associated with the receptor class were selected and further refined using homogeneous refinement followed by non-uniform refinement to yield the final reconstruction. Overall resolution was estimated using the gold-standard Fourier shell correlation (FSC) at the 0.143 criterion. Local resolution maps were calculated in cryoSPARC from the local Fourier shell correlation (FSC) between independently refined half-maps, using the mask generated during non-uniform refinement. To generate maps coloured by local resolution, the local resolution map along with the main map were opened in UCSF ChimeraX v1.10.1^71^ and processed using the surface colour tool.

### Model building, refinement, validation, analysis and presentation

The starting models were generated using AlphaFold2^72^. Iterative rounds of refinement and manual model building were performed in Coot 0.9.8.96^73^ and Phenix 2.0-5848^74^. Non-crystallographic symmetry (NCS) constraints (four groups) were applied during refinement in Phenix: chains A–B (TfR), C–D (scFv light chains), E–F (scFv heavy chains), and G–H (glycans on the two TfR molecules). The models were validated using MolProbity^75^. Structural overlays were generated using Matchmaker function in UCSF ChimeraX and C_α_ rmsd values measured using the rmsd function. Structural presentations for Fig. were produced using UCSF ChimeraX v1.10.1 or Pymol 2.1.1 (Schrödinger). Histidine mutations were introduced into the mTfR1 model using ChimeraX v1.10.1 to automatically select non-clashing common rotamers.

### Antibody production and purification for binding assays and *in vivo* testing

Expi293F suspension cells (25 ml; Thermo Fisher Scientific), grown in Expi293 Expression Medium (Gibco, A1435101) to a density of 4×10^6^ cells/ml at 37 °C, 8% CO_2_, and 130 rpm shaking, were transfected with 50 μg DNA and 200 μg PEI Max, using a 1:1 HC: LC DNA ratio for BiV-8D3 or a 1:1:1 HC:LC:Fc DNA ratio for MoV-8D3. The following day, cells were centrifuged and resuspended in 50 ml fresh medium, and the supernatant was harvested 6 days later. 8D3 antibodies were purified by nickel affinity chromatography as described above. For binding assays, antibodies were concentrated and buffer-exchanged by ultrafiltration using 50 kDa molecular weight cut-off membranes (Amicon, UFC9050). For *in vivo* testing, antibody production was scaled up to between 500-1000 ml culture media. The same initial purification process was applied but all buffers were prepared in endotoxin free water (Thermo Scientific 10307052). Antibodies for *in vivo* studies were further purified by SEC on a Superdex 200 column into sterile Dulbecco’s PBS (DPBS; Sigma-Aldrich, D8537) without Ca^2+^ or Mg^2+^. Peak fractions corresponding to 8D3 antibodies were collected and concentrated to 5-10 mg/ml. Antibodies used for *in vivo* experiments were endotoxin-depleted using endotoxin removal spin columns (Thermo Fisher Scientific, 88274), with the final endotoxin levels <50 EU/mg as determined using a Chromogenic Endotoxin Quantification Kit (Thermo Fisher Scientific, A39552). Antibodies were snap-frozen in liquid nitrogen and stored at −80 °C until needed.

### Fluorescence-based cell binding assay

bEnd3 and HEK293 cells were seeded at 5 × 10^4^ cells per well in black 96-well microplates (Greiner, 655086) pre-coated with 0.01% (w/v) poly-L-lysine (Sigma-Aldrich, P1399). Cells were cultured in Dulbecco’s Modified Eagle Medium (DMEM; Merck, D5671) supplemented with 2 mM L-glutamine (Gibco, 25030024), 1 mM non-essential amino acids (Gibco, 11140035), and 10% (v/v) fetal bovine serum (FBS; Gibco, 10270106) (hereafter referred to as complete DMEM) and incubated at 37°C, 5% CO_2_. For HEK293 stable cell line, cultures were maintained in blasticidin (2 µg/mL; Corning, 30-100-RB) and induced with doxycycline (2 µg/mL; Sigma D9891-1G) for 48 h prior to the assay. Serial dilutions of 8D3 antibodies were prepared in complete DMEM and added to cells for 2 h, either on ice or at 37°C to allow binding to reach equilibrium. Cells were washed twice with ice-cold PBS containing 0.5% (w/v) BSA (PBSB_0.5%_) and then incubated with AF488-conjugated anti-human IgG (Invitrogen, A-11013; 1:250) for 5 min on ice. After three additional washes with PBSB_0.5%_, relative fluorescence units (RFU) were measured on the Tecan Infinite 200 Pro plate reader, excitation 485 nm emission 525 nm. K_d_ values were obtained from equilibrium binding curves fitted using Graphpad Prism 10 and the equation:

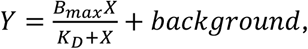

where *Y* is the measured RFU, *x* is the ligand concentration, *B_max_* is the maximal binding, and *K_d_* is the dissociation constant in the same unit as *X*. The background term represents the fluorescence signal measured in cells not expressing mTfR at the corresponding 8D3 concentrations. For the RI7 competition experiment, cells were first incubated with 50 nM RI7 (Biolegend, 113802), prior to incubation with varying concentrations of BiV-8D3 in the presence of 50 nM RI7 for 5 mins on ice, washes, and incubation with secondary as stated above.

### Internalisation assay

bEnd3 cells seeded in 96-well microplates, prepared as described above, were replaced with fresh complete DMEM and immediately incubated with BiV-8D3 (160, 10, 1 or 0.3 nM) or MoV-8D3 (260, 30, 3 or 1 nM) antibody in complete DMEM for variable durations up to 2 h at 37°C (5% CO₂). Cells were then washed twice with ice-cold PBSB_0.5%_ to arrest cell trafficking and remove unbound antibody. To quantify surface TfR cells were labelled on ice with saturating concentrations of corresponding BiV-8D3 (160 nM) or MoV-8D3 (260 nM) for 2 h. After washing, bound antibody was detected using DyLight 488-conjugated goat anti-human Fc domain antibody (Invitrogen, SA5-10134; 1:250) for 5 min on ice, followed by three washes with ice-cold PBSB_0.5%_. Fluorescence was measured on a Tecan plate reader, excitation 485 nm emission 525 nm.

### Dissociation assay

bEnd3 cells in 96-well microplates, prepared as described above, were switched to fresh complete DMEM and immediately incubated with BiV-8D3 (10 nM) or MoV-8D3 (30 nM) added to cells at staggered start times such that all conditions underwent a 2 h incubation at 37°C (5% CO₂) to reach equilibrium, and then all dissociation time points ended simultaneously. After the 2 h incubation, antibody-containing medium was removed and replaced with fresh medium, and cells were allowed to dissociate for variable durations up to 4 h at 37°C (5% CO_2_). Cells were then washed with ice-cold PBSB_0.5%_, followed by incubation with AF 488-conjugated anti-human IgG (Invitrogen, A-11013; 1:250) for 5 min on ice to detect remaining surface-bound 8D3. After three washes with PBSB_0.5%_, fluorescence was measured on a Tecan plate reader, excitation 485 nm emission 525 nm. The k_off_ values were determined by fitting dissociation curves in Graphpad Prism 10 using a one-phase exponential decay model:

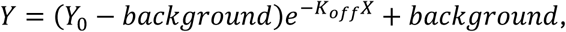

where *Y* is the measured RFU, *X* is time, *Y_0_* is the binding signal at time 0, and k_off_ is the dissociation rate constant in min^−1^.

### Flow Cytometry analysis of TfR1 surface expression

bEnd3 cells were seeded in 12-well plates at 2 × 10^5^ cells per well in complete DMEM. HEK293S mTfR FL stable cells were seeded in 24-well plates at 7.5 × 10^4^ cells per well in complete DMEM with blasticidin (2 µg/mL) and induced with doxycycline (2 µg/mL). To assess the effect of 8D3 on TfR1 surface expression, after 24 h for bEnd.3 cells and 48 h for HEK293S cells, culture medium was replaced with serial dilutions of BiV-8D3 or MoV-8D3 prepared in complete DMEM, followed by overnight incubation at 37°C. Control cells were switched into matching media lacking 8D3 to define baseline TfR1 surface staining.

After overnight treatment, cells were washed once with PBS, detached with Trypsin–EDTA, resuspended in 500 µL fresh medium, and passed through 30 µm CellTrics filters (Partec/Sysmex, 04-0042-2316) by gravity flow to reduce cell clumping. Cells were the incubated with a saturating concentration of the corresponding 8D3 (160 nM for BiV-8D3; 260 nM for MoV-8D3) for 1 h at 4°C with end-over-end rotation to label surface TfR1. Cells were washed twice with ice-cold PBS containing 0.1% (w/v) BSA and 2 mM EDTA (PBSB_0.1%_) by centrifugation (300 × g, 3 min, 4°C), then incubated with DyLight 488-conjugated anti-human Fc detection antibody (1:100; Invitrogen, SA5-10134) for 5 min on ice. Following two washes with PBSB_0.1%_ (300 × g, 3 min, 4°C), cells were fixed in 4% PFA in PBS for 5 min on ice, washed twice more, resuspended in 500 µL PBSB_0.1%_, and analyzed on a BD Accuri™ C6 Plus flow cytometer. Cells were first gated on a forward scatter area (FSC-A) versus side scatter area (SSC-A) plot to exclude debris, and singlets were then selected using forward scatter height (FSC-H) versus FSC-A. FSC/SSC gates were set according to the scatter characteristics of the main cell population and adjusted separately for bEnd3 and HEK293 cells. Fluorescence was excited with the 488 nm laser and detected in the FL1 channel (533/30 nm).

### *In vitro* immunofluorescence detection of 8D3/ TfR1

bEnd3 and HEK293 cells were seeded at 5 × 10^4^ cells per well onto 0.01% (w/v) poly-L-lysine–coated 19 mm coverslips in 12-well plates (Corning, 3513) and cultured for 48 h in complete DMEM. For HEK293 stable cell line, cultures were maintained in blasticidin (2 µg/mL) and induced with doxycycline (2 µg/mL) for 48 h prior to the assay.

To test the impact of 8D3 on TfR1 surface localisation, BiV-8D3 (160 nM) or MoV-8D3 (260 nM) was added to the culture medium at 37°C (5 % CO_2_) for 24 h before fixation. Untreated control cells received fresh medium only. Cells were washed with ice-cold PBS and fixed with 4% PFA/PBS for 10 min at room temperature, followed by two washes with PBS containing 0.5% (w/v) BSA (PBSB_0.5%_). To define maximal 8D3 staining in the no-treatment controls, fixed untreated cells were incubated with BiV-8D3 (160 nM) or MoV-8D3 (260 nM) for 1 h immediately prior to secondary detection. Bound 8D3 was detected by incubating coverslips with DyLight 488-conjugated goat anti-human Fc domain antibody (Invitrogen, SA5-10134; 1:250) for 20 min at room temperature, followed by PBS washes.

For colocalisation analysis with LAMP1, bEnd3 cells were treated with BiV-8D3 or MoV-8D3 for 1 h or 24 h as described above. For EEA1 colocalisation analysis, bEnd3 cells were treated with BiV-8D3 (160 nM), 8D3-KO (160 nM) or MoV-8D3 (260 nM) for 24 h at 37°C. For colocalisation analysis with clathrin heavy chain (CHC), bEnd3 or HEK293 cells were instead incubated with MoV-8D3 (260 nM) on ice for 1 h. Following 8D3 treatment, cells were fixed with 4% PFA/PBS for 10 min, permeabilised in PBS containing 0.1% Triton X-100 for 15 min and blocked in PBS containing 0.5% (w/v) BSA (PBSB_0.5%_) for 30 min at room temperature. Cells were then incubated for 1 h at room temperature with primary antibodies against TfR1 (H68.4, sc-65882-AF488; 1:100), EEA1 (Cell Signaling, 3288; 1:100), LAMP1 (Abcam, AB24170; 1:300), and/or CHC (Invitrogen, MA1-065; 1:200). After two PBS washes, coverslips were incubated for 20 min at room temperature with the appropriate secondary antibodies (anti-human IgG AF488, Invitrogen A-11013, 1:250; anti-human IgG AF568, Invitrogen A-21090, 1:250; anti-mouse AF568, Invitrogen A-11004, 1:250; anti-rabbit AF647, Cell Signaling 44145, 1:250), washed three times with PBS, and mounted using DAPI-containing mounting medium (Abcam, ab104139).

Images were acquired on a Leica Stellaris 5 confocal microscope using a 63× oil-immersion objective. Colocalisation was quantified in FIJI (v2.16.0) by calculating Pearson’s correlation coefficient (Pearson’s R) after background subtraction.

### pH screening assay and binding assay

5 × 10^4^ stable HEK293-mTfR cells were seeded per well into a black 384-well plate (PerkinElmer, 6007660) pre-coated with 0.01% (w/v) poly-L-lysine (Sigma-Aldrich, P1399) and incubated at 37°C, 5% CO_2_. After 24 h, cells were induced with 2 μg/mL doxycycline and incubated for a further 48 h. For the binding assay, 8D3 variants in media were diluted 4-fold in either a physiological pH buffer (1x PBS, 50 mM HEPES, 0.5% BSA, pH 7.4) or a low pH buffer (110 mM MES, 145 mM NaCl, 2.7 mM KCl, 1 mM CaCl2, 0.5 mM MgCl2, 0.5% BSA, pH 5.5) for a 5-point binding curve in a transparent reagent plate and chilled on ice. Cell media from the 384-well plate was removed and the plate placed on ice. 25 μL/well of the serial dilutions of the 8D3 variants was added and incubated at 4°C for 1 hour. After, the media was removed and two wash steps with ice cold 1x PBS were performed. Goat anti-Human IgG (H+L) Cross-Adsorbed Antibody, Alex Fluor 594 (ThermoFisher, A-11014) was diluted 1:500 in PBS + 0.5% BSA and 20 μL/well was added to the 384-well plate and incubated at 4°C for 15 minutes. After two further wash steps with ice cold 1x PBS, fluorescence was read on a Tecan microplate reader, excitation 585 emission 625 nm. For purified proteins (8D3 variants or MEM-75 or MEM-189) the binding assay was performed as in the “Fluorescence-based cell binding assay” section, except that binding curves were compared in the physiological pH buffer versus low pH buffer described here.

### Animal Experiments

All animal experiments were conducted in accordance with the Animals (Scientific Procedures) Act 1986 Amendment Regulations 2012, under UK Home Office Project Licences held by E.St.J.S. (P7EBFC1B1 and PP5814995) and approved by the University of Cambridge Animal Welfare Ethical Review Body.

Male C57B/6J mice (8–12 weeks; Envigo) were housed under conventional conditions in a temperature-controlled room (21 °C) on a 12 h light/dark cycle. Mice were weighed prior to dosing and injected intraperitoneally with 8D3 antibody variants (20 mg/kg; maximum dose volume 20 mL/kg). Antibodies were formulated in sterile DPBS, with endotoxin levels <50 EU/mg.

At 24 h post-injection, mice were terminally anaesthetised with sodium pentobarbital (200 mg/kg, i.p.) and transcardially perfused with heparinised ice-cold PBS (H3393-100KU, Sigma, 10IU/mL); for immunofluorescence experiments, perfusion was followed by 4% (w/v) paraformaldehyde (PFA) in PBS (pH 7.4). Immediately subsequent to heart ejection, blood (540 µL) was collected into syringes pre-filled with 60 µL of 3.2% sodium citrate (VWR). Brains were harvested, and for ELISA samples the olfactory bulbs and cerebellum were removed. Blood samples were centrifuged (1500 × g, 15 min, 4 °C) and plasma supernatant was collected.

### Brain sectioning and immunofluorescence

Excised brains were post-fixed overnight in 4% PFA in PBS (pH7.4) at 4°C, then cryoprotected in 30% (w/v) sucrose in PBS at 4°C until the tissue sank. Brains were embedded in Tissue-Tek O.C.T. compound and sectioned on a cryostat (Leica CM1900). Sagittal sections (40 µm) were collected onto adhesive glass slides (Epredia, J1800AMNZ). Sections were washed once in PBS containing 0.3% (v/v) Triton X-100 (PBST_0.3%_), permeabilised in PBST_0.3%_ for 1 h, and blocked for 1 h at room temperature in 10% normal goat serum (Gibco, 16210064) prepared in PBST_0.3%_. To detect injected 8D3 antibodies, AF568–conjugated anti-human secondary antibody was diluted in blocking buffer (1:1000) and incubated with sections overnight at 4°C. Sections were washed three times in PBST_0.3%_, mounted with DAPI-containing mounting medium (Abcam, ab104139), and imaged on a Leica Stellaris 5 confocal microscope. Parenchymal fluorescence was quantified in FIJI (v2.16.0) as mean intensity from five randomly selected 10 × 10 pixel regions per section placed in parenchyma while avoiding vessel-associated staining.

### ELISA analysis of antibody concentrations

Harvested brains were weighed and lysed by adding 3 volumes (v/w) of ice-cold lysis buffer (PBS containing 1% Triton X-100 and cOmplete™ protease inhibitor cocktail; Roche, 11836170001). Tissue was homogenised using a 2 cm³ glass mortar-and-pestle homogeniser (two sets of 10 clockwise strokes with 5 s rests), then incubated for 1 h at 4°C with end-over-end rotation. Homogenates were clarified by centrifugation (14,000 × g, 20 min, 4°C), and supernatants were collected for ELISA. Plasma and brain homogenate samples were pre-blocked 1:1 (v/v) with 2% (w/v) Marvel milk in PBS (MPBS_2%_) overnight at 4°C on a roller mixer. Nunc MaxiSorp 96-well plates (Thermo Fisher Scientific, 442404) were coated overnight at 4°C with anti-human IgG Fc capture antibody (Invitrogen, A18825; 1:2500). Plates were washed once with PBST (PBS + 0.1% Tween-20) and blocked with MPBS_2%_ for 1 h at room temperature. After one PBST wash, serial dilutions of pre-blocked plasma and brain homogenate (prepared in PBST) were added and incubated for 1 h at room temperature. In parallel, serial dilutions of the corresponding purified 8D3 antibody variants were prepared in PBST to generate standard curves. Plates were washed five times with PBST and incubated with anti-human IgG Fc–HRP detection antibody (Invitrogen, A18823; 1:1000 in PBST) for 30 min at room temperature. After five PBST washes, TMB substrate (Thermo Scientific, 34028) was added for 5 min, and the reaction was stopped with 2 M H₂SO₄. Absorbance was read at 450 nm. Sample concentrations were calculated from the standard curve and corrected for dilutions. 8D3 concentrations in brain were additionally normalised by correcting for the homogenisation dilution (3× lysis buffer relative to tissue weight). Receptor occupancy was calculated using K_d_ values measured on bEnd3 cells at 37°C for BiV-8D3 and MoV-8D3, and on HEK293 mTfR FL+ stable cells at 4°C for the other 8D3 variants.

### Statistical analysis

All statistical analyses were performed in GraphPad Prism (v10.0). Two-group comparisons were analysed using a two-tailed unpaired t-test with Welch’s correction (α = 0.05). Comparisons among multiple groups were analysed by one-way ANOVA followed by Tukey’s multiple-comparisons test (α = 0.05). Data are presented as mean ± SEM unless otherwise stated.

