## Supplementary material for "Molecular mechanism of action of a blood brain barrier shuttle antibody": Supp Figs 1-7 & Tables 1-3

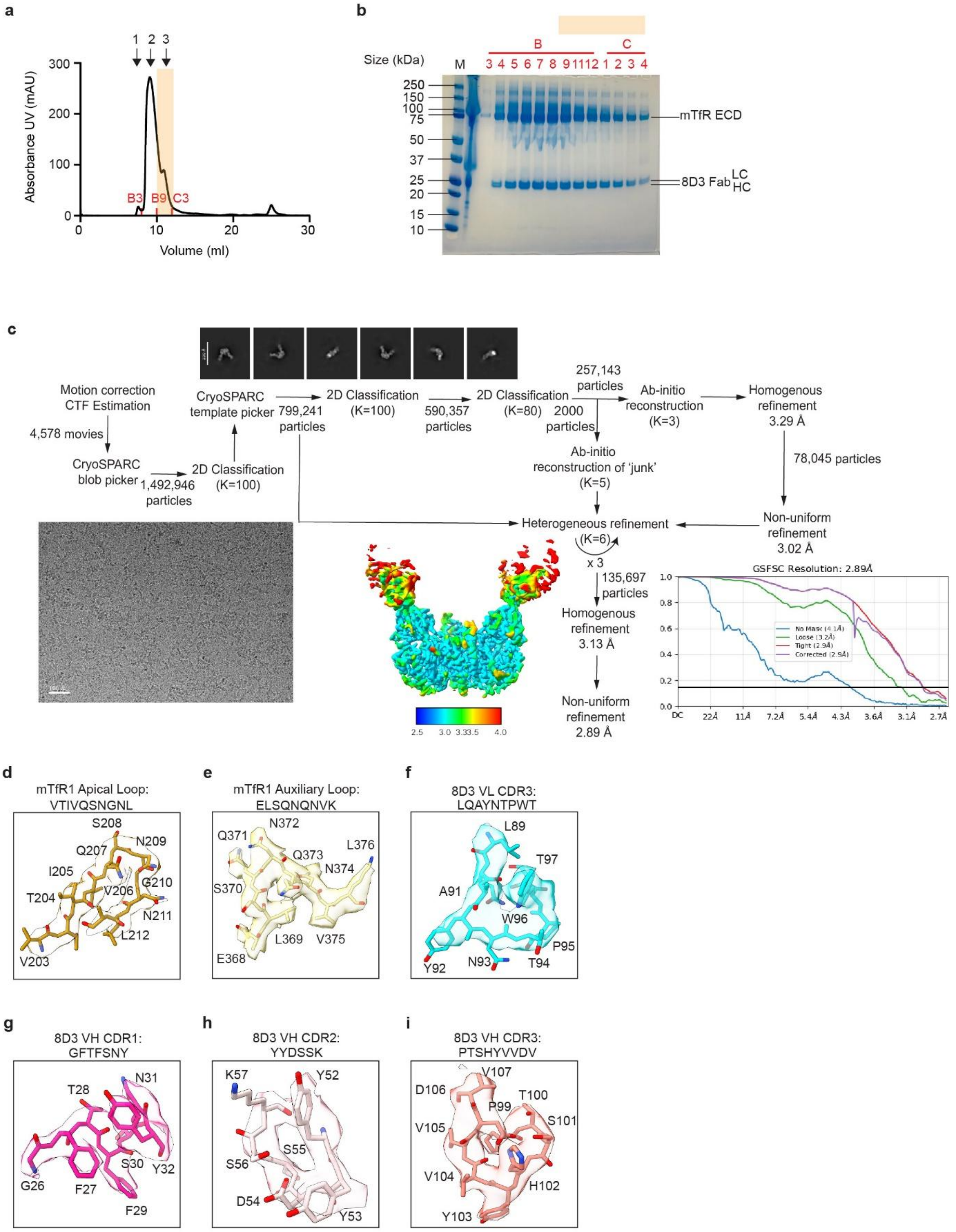

**Extended Data Fig. 1. mTfR1 ECD purification, cryo-EM data processing, binding loop electron density. a,** Size-exclusion chromatography (SEC) trace for mTfR1-8D3 complex run on a Superose-6 10/300GL column. Purified mTfR1 protein eluted as a broad oligomeric peak by size exclusion chromatography and precipitated if stored overnight. 1, aggregate peak in void volume; 2, oligomeric peak; 3, anticipated dimer peak; selected fractions indicated in red; peach highlight region indicates fraction selected for cryo-EM. **b**, Coomassie stained reducing SDS-PAGE gel of SEC fractions (5 µl loads). **c**, Image processing workflow for mTfR1 ECD in complex with brain shuttle Fab 8D3. Cryo-EM of a peak containing a presumed dimeric species resulted in 2D classes of a V-shaped protein dimers. Movie frames were motion-corrected and contrast transfer function (CTF) parameters estimated in RELION. Subsequent processing was performed in cryoSPARC, including particle picking, 2D classification, and ab initio 3D reconstruction. The most well-resolved classes were refined to generate an initial reference map, which was used for multiple rounds of heterogeneous refinement alongside low-quality classes using the full particle stack. The largest particle subset was selected for further refinement, yielding the final reconstruction. The final map was refined without imposed symmetry (C1) to a global resolution of 2.89 Å (FSC 0.143 criterion), with local resolution estimates shown. **d-i**, Electron density, protein sequence and model build for **d**, mTfR1 main apical loop, **e**, mTfR1 auxiliary apical loop **f**, 8D3 $V_L$ CDR3, **g**, 8D3 $V_H$ CDR1, **h**, 8D3 $V_H$ CDR2, **i**, 8D3 $V_H$ CDR3. Nitrogen atoms – blue, oxygen atoms – red.

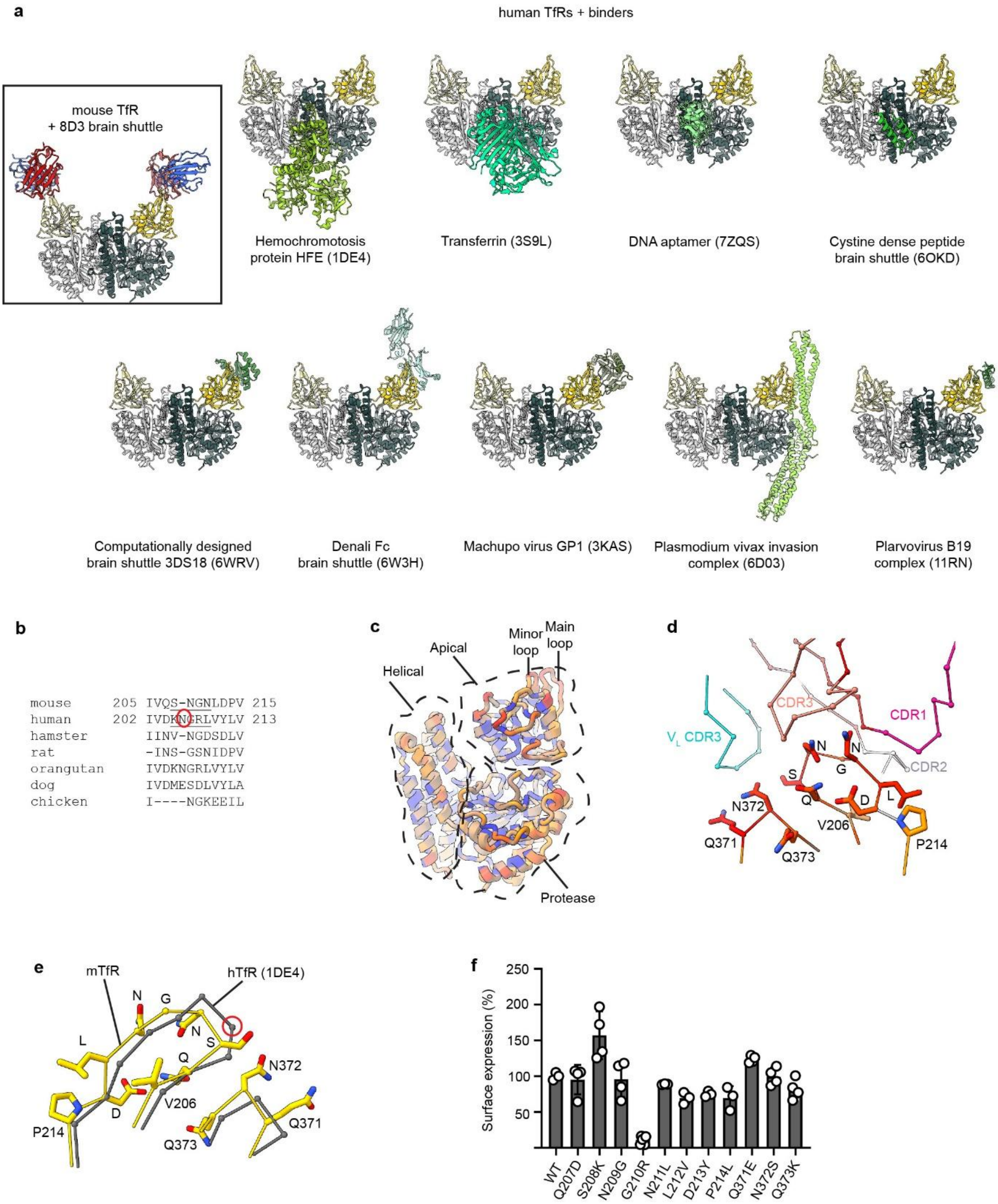

**Extended Data Fig. 2. mTfR versus hTfR1 binder structures. a**, Comparison binder locations on the mTfR1-8D3 complex (boxed) versus previously published hTfR1-binder complexes (ribbon representation). TfR1 Protease+helical domain, dark/light grey; hTfR1 apical domain, yellow/pale yellow. Binder colors are: 8D3 Fab $V_H$ domain, blue; 8D3 Fab $V_L$ domain, red; other protein binders in varying shades of green/grey/blue. Top panel shows binders that bind to the hTfR1 helical domain. Lower panel shows binders that bind to the hTfR1 apical domain. Corresponding Protein Data Bank (PDB) identification codes are provided in parentheses. **b**, TfR1 main apical loop protein sequence alignment. An additional hTfR1 Asn residue versus mTfR1 is indicated by red circle. **c**, Ribbon representation of TfR1 colored by sequence conservation (red, most conserved; blue, least conserved). Both the labelled apical loops targeted by 8D3 are red/orange. **d**, Closeup $C_\alpha$ representation colored as in (D) with apical loop side chains shown. **e**, Structural overlay of mTfR1 versus hTfR1 (1DE4) main apical loop showing similar $C_\alpha$ trajectory. hTfR1 Asn insertion indicated by red circle. **f**, Histogram showing surface expression of mTfR1 wild-type (WT) and single residue substitutions in the main apical loop (207-214) or alternate apical loop (371-373). Surface expression measured from adherent cells seeded in 96-well plates stained using streptavidin-AlexaFluor488 against a C-terminal (extracellular) twin strep tag on each mTfR1 subunit of the dimeric receptor. Values are mean ± SEM from *n* = 3-4 independent experiments.

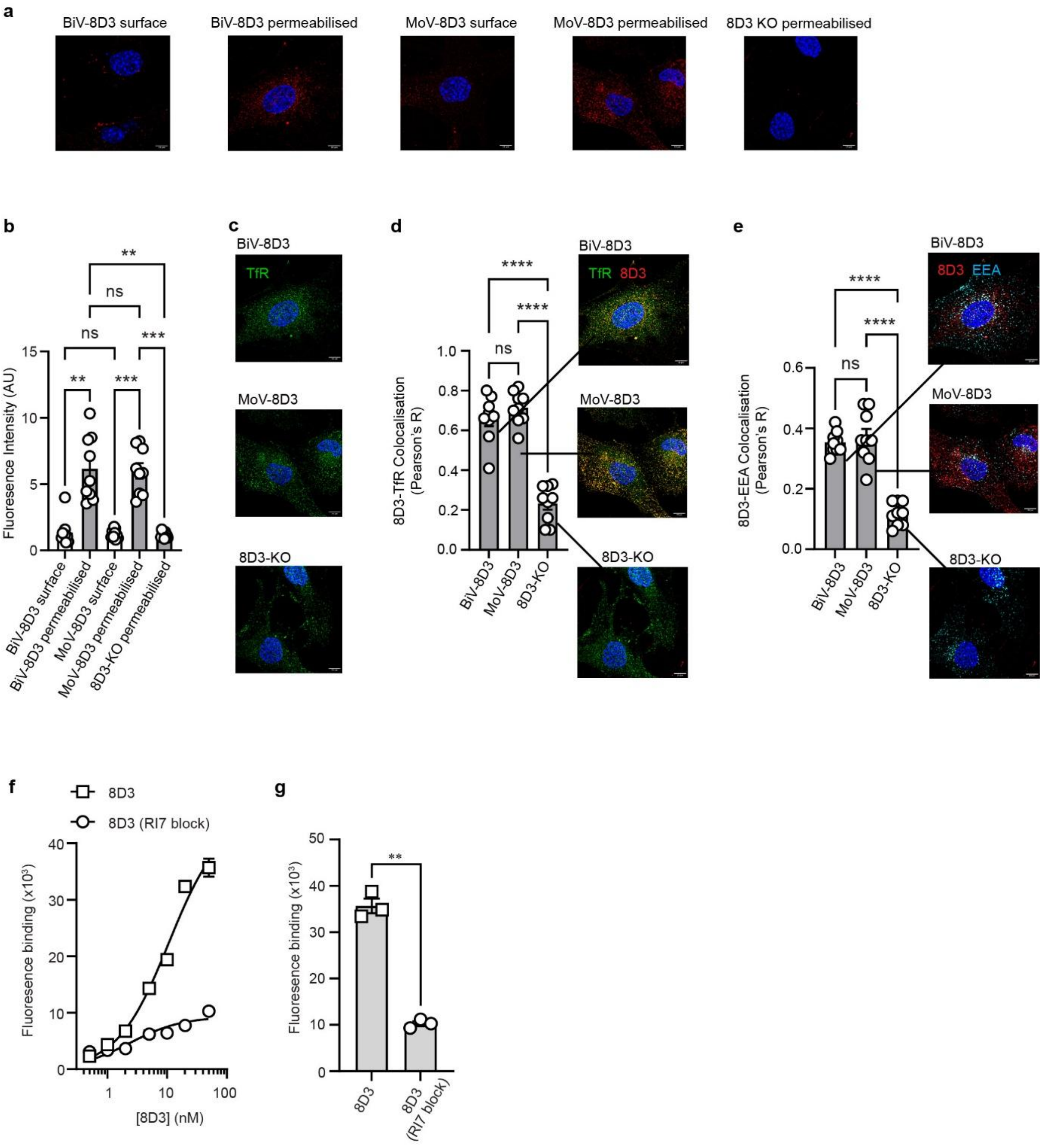

**Extended Data Fig. 3. Impact of 8D3 on intracellular mTfR1 distribution and competition with RI7. a**, 8D3 fluorescence intensity and distribution after 24 h incubation at 37 °C with bEND3 cells for BiV-8D3, MoV-8D3 and 8D3-KO, comparing surface versus permeabilised staining. **b**, Cell intensity quantification from, **a**. **c**, Permeabilised cell mTfR1 distribution using anti-mTfR1 (H68.1). **d-c**, Quantification of **d**, mTfR1–8D3 and **e**, 8D3-EEA1 colocalization in permeabilised cells, expressed as Pearson's correlation coefficient following incubation with BiV-8D3, MoV-8D3, or 8D3-KO at 37°C for 24 h, along with representative images for each treatment. **b, d, e,** Values are mean ± SEM determined from are *n* = 9 individual cells from 3 independent experiments. **f-g**, Fluorescence-based on-cell binding assay measuring 8D3 binding in the absence or presence of pre- and co-incubation with 50 nM RI7, to mTfR1-expressing HEK293S stable cells at 4 °C. **f**, shows dose response curves. **g**, shows binding for highest 8D3 dose, 50 nM. RI7 competes for the same epitope to block 8D3 binding. Values are mean ± SEM determined from *n* = 3 independent experiments. **, *P* < 0.01, two-tailed unpaired t-test.

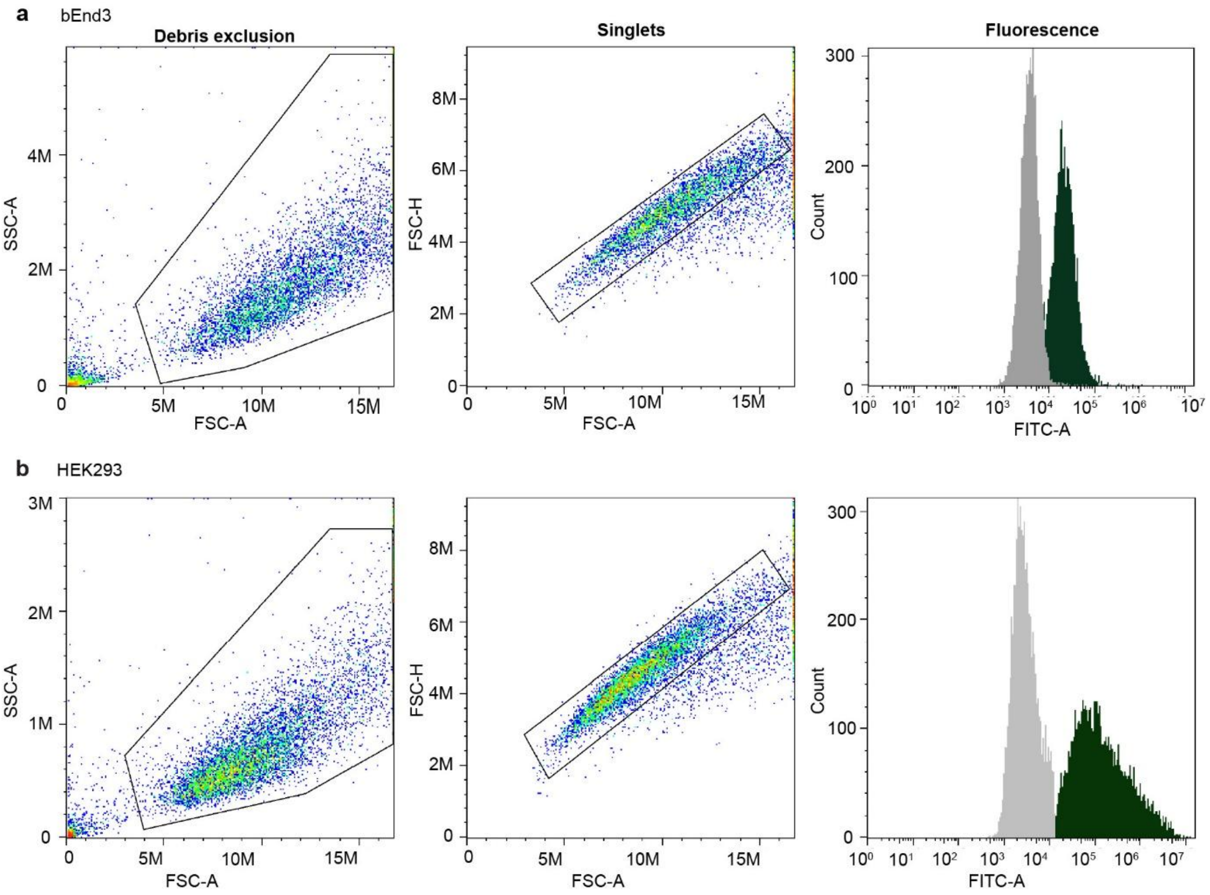

**Extended Data Fig. 4. mTfR1 surface expression by flow cytometry, bEND3 versus HEK293 cells. a-b,** Raw example flow cytometry data showing the gating strategy for **a**, bEnd3 cells and **b**, HEK cells. Representative flow cytometry plots showing sequential exclusion of debris (left panels) and selection of singlets (middle panels) before fluorescence quantification (right panels). Debris was excluded using forward scatter area (FSC-A) versus side scatter area (SSC-A), followed by singlet gating based on FSC-A versus forward scatter height (FSC-H). Right panels show representative FITC-A fluorescence histograms of the final gated singlet populations for the negative control (grey) and cells labelled with 8D3 to detect total surface mTfR1 (dark green). Raw data from *n* = 1 representative experiment shown.

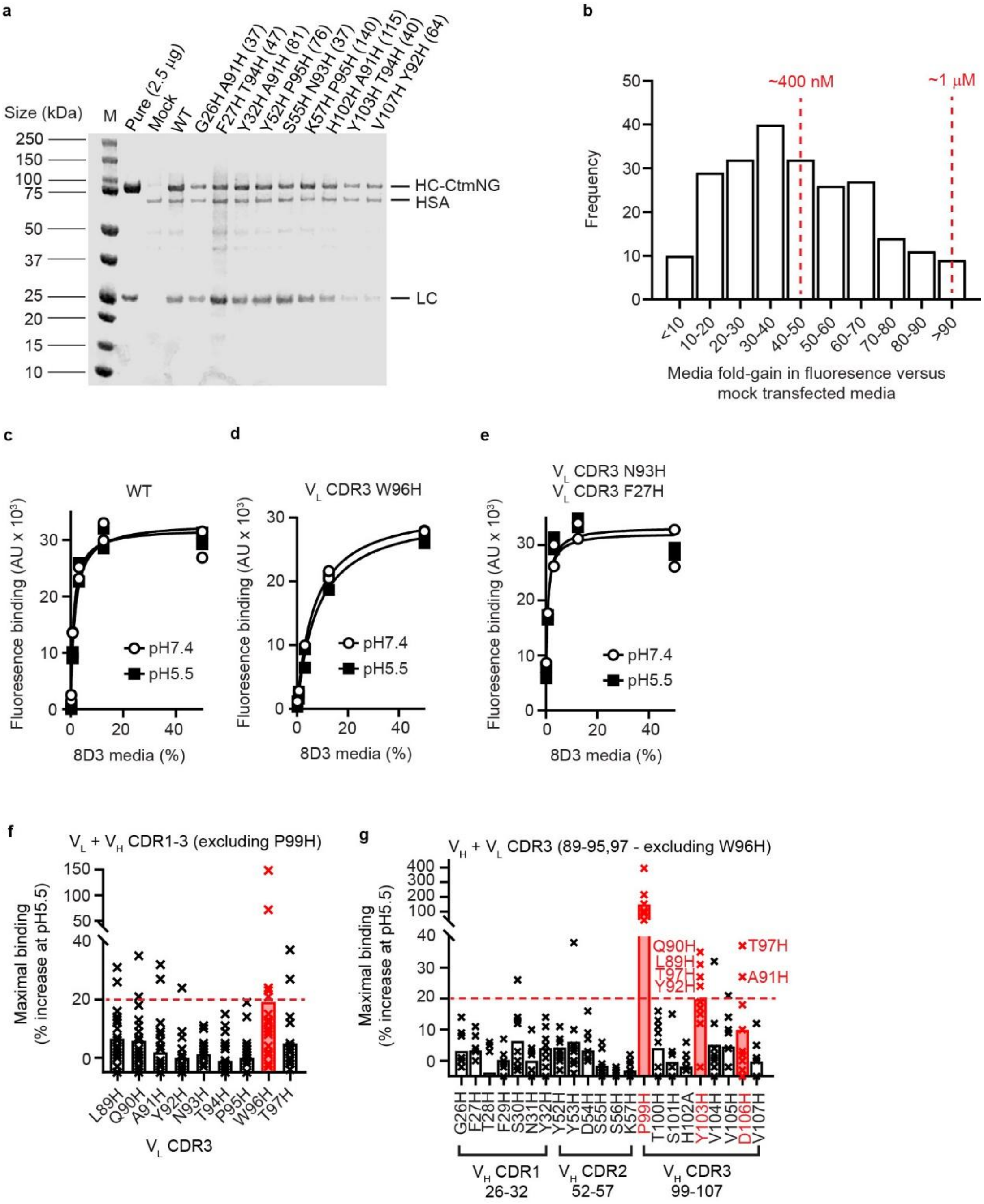

**Extended Data Fig. 5. pH-sensitive 8D3 library production and screening. a**, Coomassie stained reducing SDS-PAGE gel of Expi293F cell expression media (10 µl loads) for selected 8D3 histidine mutants. HSA, human serum albumen; mock, mock transfected cells negative control; CtmNG, C-terminal mNeon Green; pure, purified wild-type 8D3-CtmNG. **b**, Distribution of expression yields across the pH-sensitive library, plotted as fold-gain in fluorescence (from CtmNG) in expression media relative to mock-transfected media; red dashed lines indicate approximate antibody concentrations (~400 nM and ~1 µM) determined from titrations into conditioned media by known amounts of purified wild-type 8D3-CtmNG. **c-e**, Representative on-cell binding curves at pH 7.4 versus pH 5.5 using unpurified 8D3 in expression medium for **c**, wild-type BiV-8D3 and representative **d**, single and **e**, double histidine variants. **f-g**, Percent increase in maximal binding at pH 5.5 versus pH 7.4 for combined $V_L$+$V_H$ histidine variants. **f**, Arranged by each specific $V_L$ mutation combined with any $V_H$ mutation. **g**, Arranged by each specific V mutation combined with any $V_H$ mutation. Red bars indicate specific mutations where pairings reached mean increases > 10 %. Dashed line indicates a 20% increase threshold, and red highlights variants exceeding this threshold. In all experiments screening was done as *n* = 1 experiment (technical duplicates).

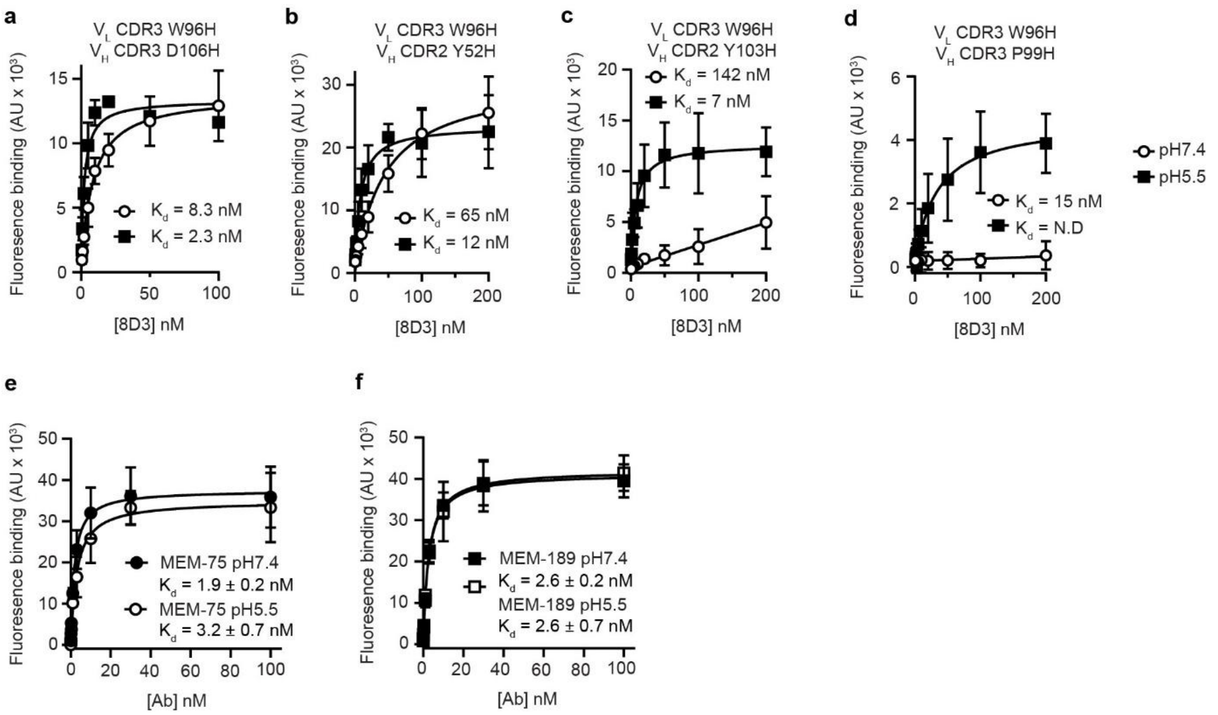

**Extended Data Fig. 6. pH-sensitive variants. a-d**, On-cell binding curves from selected purified pH-sensitive 8D3 variants carrying histidine substitutions show preferential mTfR1 binding at pH 5.5 versus pH 7.4. Values are mean ± SEM determined from binding curves from *n* = 6 independent experiments (except 8D3 $V_L$ W96H $V_H$ Y103H pH7.4, *n* = 4). **e-f**, MEM-75 and MEM-189 show marginal or no pH-sensitive binding to hTfR1. Values are mean ± SEM determined from binding curves from *n* = 5 independent experiments.

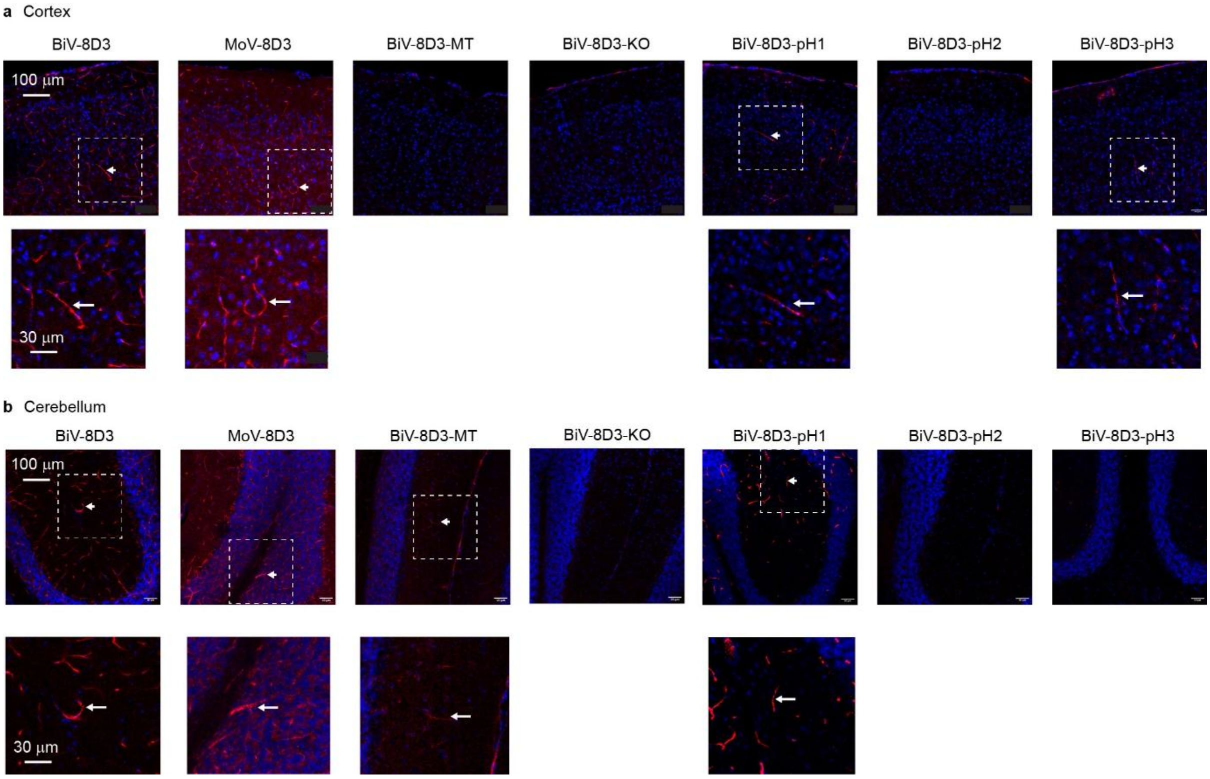

**Extended Data Fig. 7. Mouse *in vivo* receptor labelling in cortex and cerebellum. a-b**, Representative sagittal brain sections showing distribution of 8D3 antibodies in **a**, cortex and **b**, cerebellum at 24 h after intraperitoneal injection (20 mg/kg). 8D3, red (anti-human IgG staining); nuclei, blue (DAPI). Dashed boxes indicate regions enlarged below; arrows mark vessel staining.

**Supplementary Table 1. Cryo-EM data collection, processing, refinement and validation statistics.**

| | **TfR–scFv 8D3** Version 77 (EMD-57148) (PDB 29GF) |
|---|---|
| **Data collection and processing** | |
| Microscope | Titan Krios |
| Detector | Gatan K3 |
| Magnification | 130K |
| Energy filter slit width (eV) | 20 |
| Voltage (kV) | 300 |
| Exposure Time (s) | 1.61 |
| Number of Frame | 120 |
| Electron exposure ($e^-/Å^2$) | 48.02 |
| Defocus range (μm) | -2.5, -2.2, -1.9, -1.6, -1.3, -1.0, -0.8 |
| Pixel size (Å) | 0.652 |
| Symmetry imposed | *C*1 |
| Number of movies collected | 4578 |
| Initial particle images (No.) | 1,492,946 |
| Final particle images (No.) | 135,697 |
| Box Size | 416 |
| Map resolution at FSC=0.143 (Å) | 2.89 |
| Map Sharpening B Factor ($A^2$) | 79.34 |
| **Refinement** | |
| Model composition | |
| Atoms (hydrogens) | 27190 (13490) |
| Protein residues | 1724 |
| Protein atoms | 26976 (13388) |
| Glycan residues | 8 |
| Glycan atoms (hydrogens) | 214 (102) |
| Mean B-factors ($Å^2$) | |
| Protein | 112.57 |
| Glycan | 102.44 |
| R.m.s. deviations | |
| Bond lengths (Å) | 0.004 |
| Bond angles (°) | 0.635 |
| Ramachandran plot | |
| Favoured (%) | 95.56 |
| Allowed (%) | 4.44 |
| Disallowed (%) | 0.00 |
| MolProbity | |
| Overall score | 1.45 |
| Clash score | 3.27 |
| Rotamer outliers (%) | 1.08 |

R.m.s. deviations: root mean square deviations from ideal geometry.

| 8D3 chain | | pH7.4 $K_d$ (nM) | | | | pH5.5 $K_d$ (nM) | | | | |
|---|---|---|---|---|---|---|---|---|---|---|
| $V_H$ | $V_L$ | mean | S.E.M | n | Fold vs WT | mean | S.E.M | n | Fold vs WT | pH5.5 selectivity |
| WT | WT | 1.6 | 0.1 | 28 | | 1.6 | 0.2 | 28 | | |
| W96H | WT | 2.0 | 0.1 | 6 | 1 | 1.9 | 0.1 | 6 | 1 | None |
| WT | Y52H | 3.0 | 0.2 | 6 | 2 | 3.3 | 0.2 | 6 | 2 | None |
| WT | P99H | >200 | - | 6 | - | 43 | 13 | 6 | 27 | > 5-fold |
| WT | Y103H | 1.8 | 0.1 | 6 | 1 | 1.7 | 0.1 | 6 | 1 | None |
| WT | D106H | 1.7 | 0.1 | 6 | 1 | 1.6 | 0.1 | 6 | 1 | None |
| W96H | Y52H | 65 | 25 | 6 | 40 | 12 | 5 | 6 | 8 | 4-fold |
| W96H | P99H | >200 | - | 6 | | 15 | 5 | 6 | 9 | >>13-fold |
| W96H | Y103H | 142 | 16 | 4 | 90 | 7.0 | 2.2 | 6 | 4 | 20-fold |
| W96H | D106H | 8.3 | 0.3 | 6 | 1 | 2.3 | 0.3 | 6 | 5 | 4-fold |

**Table S2**

| | 8D3 construct | | | | | | |
|---|---|---|---|---|---|---|---|
| | BiV | MoV | MT | KO | pH1 | pH2 | pH3 |
| | Concentration (nM) | | | | | | |
| Plasma | 110 ± 40 | 33 ± 12 | 110 ± 20 | 430 ± 90 | 240 ± 60 | 260 ± 60 | 140 ± 30 |
| Brain | 2.4 ± 0.4 | 15 ± 3 | 3.4 ± 0.7 | 0.9 ± 0.2 | 0.5 ± 0.1 | 1.0 ± 0.3 | 1.1 ± 0.2 |
| | Predicted occupancy (%) (based on $K_D$ and plasma/brain concentration) | | | | | | |
| Plasma | 97 ± 1 | 67 ± 8 | 47 ± 4 | | 62 ± 8 | | |
| Brain | 71 ± 3 | 58 ± 5 | 3 ± 0.6 | | 1.7 ± 0.4 | | |

**Table S3**
